# Identifying novel interactions of the colon-cancer related APC protein with Wnt-pathway nuclear transcription factors

**DOI:** 10.1101/2022.06.29.498083

**Authors:** Nayra M. Al-Thani, Stephanie Schaefer-Ramadan, Jovana Aleksic, Yasmin A. Mohamoud, Joel A. Malek

## Abstract

**Background:** Colon cancer is often driven by mutations of the adenomatous polyposis coli (APC) gene, an essential tumor suppressor gene of the Wnt β-catenin signaling pathway. APC and its interactions in the cytoplasm have been well studied, however various groups have also observed its presence in the nucleus. Identifying novel interactions of APC in the Wnt pathway will provide an opportunity to better understand the nuclear role of APC and ultimately identify potential cancer treatment targets.

**Methods:** We used the all-vs-all sequencing (AVA-Seq) method to interrogate the interactome of protein fragments spanning most of the 60 Wnt β-catenin pathway proteins. Using protein fragments identified the interacting regions between the proteins with more resolution than a full-length protein approach. Pull-down assays were used to validate a subset of these interactions.

**Results:** 74 known and 703 novel Wnt β-catenin pathway protein-protein interactions were recovered in this study. There were 8 known and 31 novel APC protein-protein interactions. Novel interactions of APC and nuclear transcription factors TCF7, JUN, FOSL1, and SOX17 were particularly interesting and confirmed in validation assays.

**Conclusion:** Based on our findings of novel interactions between APC and transcription factors and previous evidence of APC localizing to the nucleus, we suggest APC may compete and repress CTNNB1. This would occur through the binding of the transcription factors (JUN, FOSL1, TCF7) to regulate the Wnt signaling pathway including through enhanced marking of CTNNB1 for degradation in the nucleus by APC binding with SOX17. Additional novel Wnt β-catenin pathway protein-protein interactions from this study could lead researchers to novel drug designs for cancer.

## Introduction

The latest statistics from the American Cancer Society show that 106,180 new colorectal cancer (CRC) cases and 52,580 deaths are expected in 2022 [1]. Most CRCs (∼80%) have mutations in the adenomatous polyposis coli (APC) gene [2, 3], which is an important regulator of the Wnt signaling pathway (Fig. 1). There are two major Wnt pathways; the first is the non-canonical signaling pathway, including Wnt/calcium and planar cell polarity (PCP) pathways, which are not CTNNB1 dependent [4]. The Wnt/calcium pathway regulates calcium influx from the endoplasmic reticulum to the extracellular space, which is essential for cellular development [4]. The second is the canonical-Wnt signaling pathway, where the function depends on the destruction complex proteins (APC, AXIN1, GSK3β, and CKI α). The deactivation of the destruction complex leads to β-catenin (CTNNB1) accumulation in the cytoplasm and its translocation to the nucleus (Fig. 1). CTNNB1 then drives gene expression leading to cell proliferation [5].

**Figure 1.**
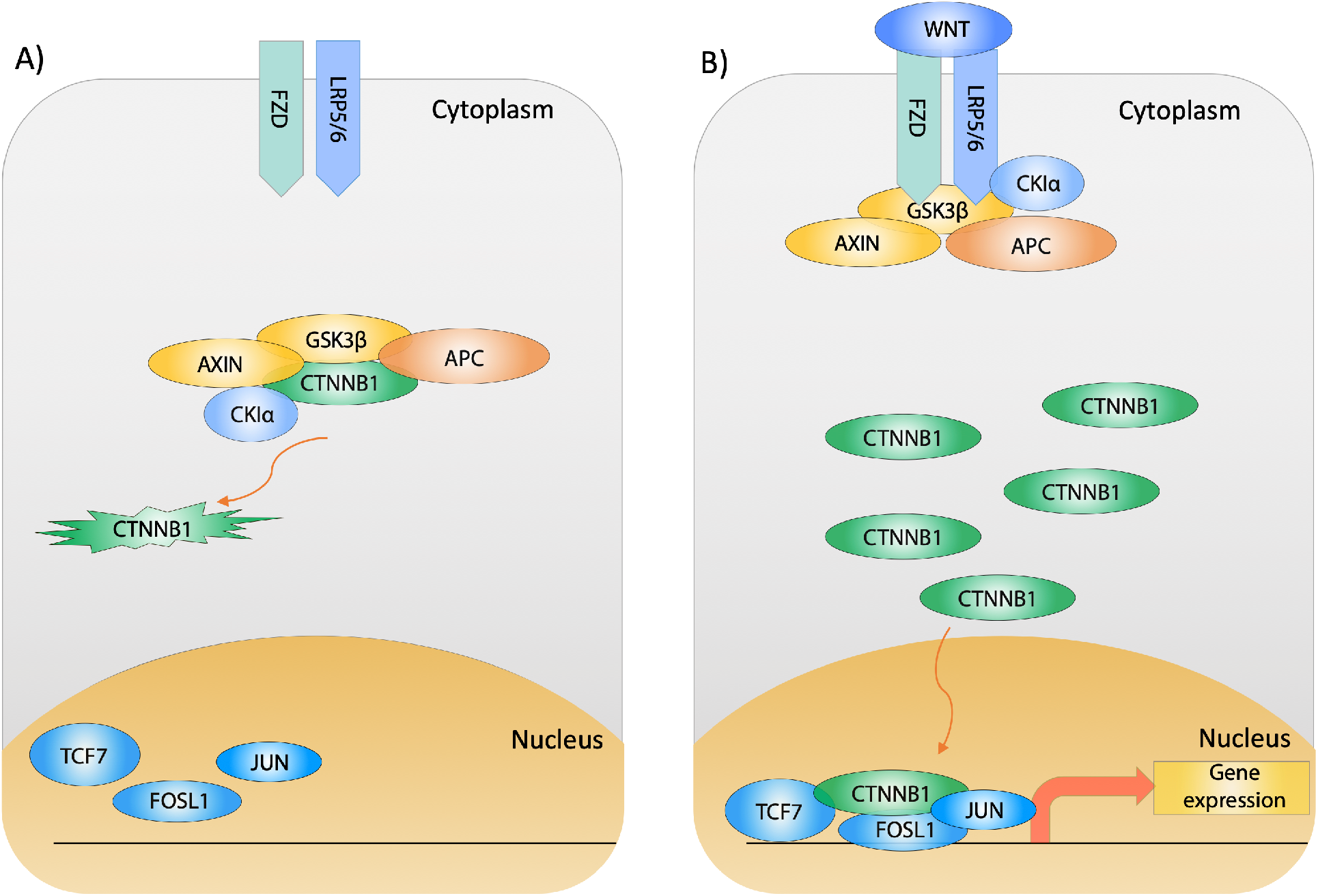
The Canonical-Wnt signaling pathway. **A)** In the absence of Wnt, the destruction complex (APC, AXIN, CK1α, and GSK3β) binds and phosphorylates CTNNB1 to mark it for proteasomal degradation. The reduced cytoplasmic level of CTNNB1 leads to the inactivation of Wnt target transcription factors (TCF7, FOSL1, and JUN). **B)** Upon Wnt binding, the destruction complex proteins (APC, AXIN, CK1α, and GSK3β) are recruited to bind FZD and LRP5/6 transmembrane proteins. This will lead to the accumulation of CTNNB1 in the cytoplasm and its translocation to the nucleus. Subsequently, nuclear CTNNB1 binds to Wnt target transcription factors (TCF7, FOSL1, and JUN), leading to gene expression and cell proliferation [5].

There is clinical evidence of APC mutations in prostate, breast, and gastric cancer and glioblastoma [5, 6]. Mutations in the Wnt β-catenin signaling pathway genes cause cell proliferation and uncontrolled growth [5, 7]. Tumor formation could be initiated upon a loss-of-function (LOF) mutation of APC or a gain-of-function (GOF) mutation of CTNNB1 [5], leading to gene expression, proliferation, and cell cycle progression in the absence of Wnt.

The binding of CTNNB1 with AP-1 transcription factors is associated with tumor malignancy [8]. The AP-1 transcription factors have a role in regulating the cell cycle progression. Both c-JUN (JUN) and Fra-1 (FOSL1), which are part of AP-1 transcription factors, are involved in cell proliferation and regulation of cellular differentiation [8-10]. Moreover, these transcription factors (JUN, FOSL1), along with transcription factor 7 (TCF7), drive the epithelial-mesenchymal transition and metastasis when expressed [11-13]. TCF7 is part of TCF/LEF family proteins which includes TCF7 (TCF1), LEF1, TCF7L1 (TCF3), and TCF7L2 (TCF4) [14]. The TCFs and LEF proteins are the final components needed for Wnt β-catenin signaling pathway activation [5]. The TCF proteins contain the HMG box, which is required to stabilize protein binding with DNA and induce gene expression [14]. TCF7 gene knock-out in CRC cell lines, slows tumor growth [15], indicating its importance in cell proliferation. CTNNB1 binds directly to both c-JUN and c-FOS *in vitro* and *in vivo*, showing direct protein-protein interactions (PPIs) and not just a transcriptional activation through DNA binding [8]. Additionally, there is evidence of c-JUN binding to TCF4 upon phosphorylation [15] which drives gene expression and cell proliferation in CRC tumors [8, 16]. Similarly, SRY-box transcription factor 17 (SOX17) has been shown to act as a tumor suppresser by binding CTNNB1/TCF3 or TCF4 in a complex and repressing their activity [17, 18].

APC localizes to the cytoplasm and nucleus [2, 3, 23]. However, less is known about the role of nuclear APC. We wondered if APC had a more substantial role in the nucleus than previously thought. The combination of nuclear APC and the importance of transcription factors in the WNT pathway offer an interesting axis of investigation.

To that end, we selected 60 canonical Wnt β-catenin signaling pathway genes for protein-protein interaction screening. We applied the all-vs-all sequencing (AVA-Seq) method to determine the interaction network of this signaling pathway with the intent to identify known and novel interactions between the 60 proteins with the added feature of localizing which regions of the protein are involved in the interactions. It is well known that various protein-protein interaction methods do not recover all interactions. So even well-studied pathways such as the WNT pathway would benefit from analysis with new methods [19, 20]. The all-vs-all sequencing (AVA-Seq) method is a novel approach for detecting PPIs. It is based on the bacterial two-hybrid system of Dove and Hochschild with several significant changes [21]. AVA-Seq allows proteins of interest to be pooled together and fused to the DNA-binding domain (DBD) and the transcription activation domain (AD) on a single plasmid. If the tested proteins interact, it leads to the expression of the *HIS3* gene and the cell’s survival in histidine dropout media [22]. An interaction is reported when there is a significant growth in the presence of a competitive inhibitor of histidine, 3-amino-1,2,4-triazole (3-AT), compared to the control sample. The growth of cells in the presence of 3-AT indicates a protein interaction. Interactions are quantified by increases in the frequency of cells harboring the interacting fragments in liquid culture versus other cells in the same pool. This is evidenced by next-generation sequencing read count increases.

Our method recovered 74 known interactions for which there is strong evidence in the literature and 703 novel PPIs. Of particular interest were novel interactions between APC and nuclear transcription factors. Namely APC with SOX17, TCF7, JUN, and FOSL1. Several of these interactions were subjected to a secondary validation using full-length and fragmented proteins. Finally, we present potential implications from the interactome data.

## Materials and Methods

### Amplification of human clones

60 Human ORF clones were purchased from GenScript (https://www.genscript.com) for the Wnt pathway (Table S1). Clones were PCR amplified using T7 forward (5’-TAA TAC GAC TCA CTA TAG GG -3’) and BGH reverse (5’-TAG AAG GCA CAG TCG AGG -3’) primers from the pcDNA3.1+/-C-(K)-D vector using standard methods (NEB 2x Q5) which include 5-10 ng DNA per reaction, 57 °C annealing temperature and 5% DMSO for GC rich PCR products. After successful amplification, reactions were column cleaned using GenElute PCR Cleanup Kit.

### Open reading frame filtering

The 60 clones were aliquoted into 30 nM final concentration, and samples were dried and resuspended into 5 *µ*L water. A 30 nM pool was prepared by taking 2 *µ*L from each sample. The final volume of 120 *µ*L is split into two reactions, each with 50 *µ*L, and sheared using a Covaris focused-ultrasonicator. Sheared DNA was end-repaired (NEB E6050S) followed by Ampure cleaning and ligation (NEB M0202S) into pBORF filtering vectors, which were described previously [22, 24].

WNT pathway selected open reading frames (ORFs) in pAVA were transformed into NEB Turbo (NEB C2986; discontinued) cells to obtain more than 20 million colonies split into two libraries of approximately 10 million each. DNA was extracted and quantified using Qubit HS. 2 ng DNA was transformed into the Validation Reporter (VR) strain (Agilent Technologies #200192; discontinued) to obtain 30-40 million transformants using electroporation. As described previously, the 3-amino-1,2,4-triazole (3-AT) selection is performed [22]. Fragment pairs grown in the absence of 3-AT (0 mM conditions) serve as a baseline for the number of read counts. The experiments had three replicates for each condition (0 mM, 2 mM, 5 mM 3-AT), resulting in 9 samples. This was repeated, resulting in two separate transformation events to maximize the screening area (meaning two replicates of 9 samples).

### Data Analysis

Data analysis was performed as previously [24]. Briefly, raw sequencing data of the plasmids containing the paired fragments grown in selective media were translated and aligned to the Wnt ORF clone database using DIAMOND Blastp [25]. In the AVA-seq method, paired-end reads reveal which two protein fragments were tested against each other. Statistically significant increases in the frequency of a pair of fragments in selective media over non-selective media indicate higher growth and likely a protein interaction between the two fragments. After Blastp analysis, paired-end read counts were normalized and tested for statistically significant increases using EdgeR. An interaction was called with a log2 fold-change (Log_2_FC) of 1.5 or greater, and we allowed a false-discovery rate (FDR) with multiple testing adjusted *p*-value of less than 5% (0.05).

### Protein Expression

Specific DNA fragments were ordered from TWIST Bioscience and optimized for *E. coli* expression (Table S3) except for SOX17. SOX17 fragments were PCR amplified using primers containing Electra cloning sites. SOX17^FL^ primers: (5’-ATG AGC AGC CCG GAT GC -3’) and (5’-TCA CAC GTC AGG ATA GTT GCA GTA -3’); SOX17^88^: (5’-ATG CAG CAG AAT CCA GAC CTG -3’) and (5’-CAG GAG GCC CGG AAT -3’); and SOX17^216^: (5’-ATG GGC TAC CCG TTG CCC AC -3’) and (5’-TCA CAC GTC AGG ATA GTT GCA GTA -3’). TWIST fragments and SOX17 (amplified PCR products) were ligated into bacterial expression vector pD454 plasmids (pD454-MBP or pD454-GST) using Electra reagents kit (atum.bio EKT-03) following ATUM Bio-protocol. The ligation was directly transformed into NEB-5-alpha electrocompetent cells (NEB C2987H) and plated on LB-agar supplemented with carbenicillin (100 µg/mL). DNA was extracted, and constructs were sequence confirmed, followed by transformation into BL21 DE3 chemi-competent cells for protein expression. 10 mL of overnight culture were used to inoculate 1 L of fresh LB media supplemented with carbenicillin (100 µg/mL). When the cells reached an OD_600_ of 0.5-0.6, the 3-hour expression at 37 °C was induced with 0.01-0.05 mM IPTG. The expression of the full-length or protein fragments was confirmed via SDS-PAGE (Fig. S1) and (Fig. S2).

All APC constructs contained an N-terminal GST protein, and the transcription factors had an N-terminal MBP to maximize the protein solubility [26].

### Pull-down

Cell pellets of 50 mL were used for protein expression in the pull-down experiments. Cell expressing protein fragments with an MBP tag were lysed first using 1 mL Bacterial Protein Extraction Reagent (B-PER; Thermo Scientific Catalog number: 78243) and 1-2x of protease inhibitor (Thermo Scientific A32963). Overexpressed MBP without a fusion protein was lysed and used as a negative control. For the negative control MBP, twice the amount is used for pull-down compared to the tested fragments of Wnt transcription factors. Bacterial cell lysis is incubated for 15 minutes at room temperature, followed by 4 °C centrifugation at 14,000 rpm for 10 minutes. 100 µL of amylose resin (NEB E8021S) is equilibrated with 1x TBS. The soluble fraction of the cell lysis is added to the pre-equilibrated resin and incubated for 1 hr rotating at 4 °C. Approximately 40 minutes later, GST-tagged proteins are lysed using the same method as above. After 1 hr incubation of the amylose resin with MBP tagged protein, the mixture is centrifuged at 1,000 rpm for 5 minutes at 4 °C. The supernatant is carefully discarded. The MBP proteins bound to resin are gently washed twice with 1 mL 1x TBS and centrifuged. The soluble fraction of the GST-tagged proteins is added to the washed resin (MBP-tagged proteins already bound) and incubated for 2 hrs rotating at 4 °C.

Following the 2 hr incubation, the resin and protein mixtures were added to a micro Bio-spin column (Bio-Rad 7326202). The flow-through was discarded, and the resin was washed four times with 1 mL 1x TBS. The proteins were eluted from the resin using 50 µL of 1x TBS supplemented with 10 mM maltose. 10 µL of each pull-down sample was run on a reducing 12% SDS gel, and the gel was transferred to PVDF membrane for Western blot and imaged using LiCor.

Anti-GST antibody (Abcam EPR4236; 1:1,000) and anti-MBP antibody (NEB E8032L; 1:10,000) in 1x TBST supplemented with 5% low-fat milk were incubated either at 4 °C overnight or 1 hr at room temperature. The secondary antibodies, IRDye 680RD anti-mouse (LiCor 926-68070; 1:15,000) and IRDye 800CW anti-rabbit (LiCor 926-32211; 1:15,000), are compatible with the LiCor imaging system.

## Results

### AVA-Seq method applied to Wnt-signaling pathway proteins

Protein fragments from 60 Wnt pathway genes (Table S1) were enriched for codon frame 1 using an open reading frame (ORF) filtering method (see methods). The ORF filtering process enriched fragments for frame 1 by 75% and 80% for DBD- and AD-associated fragments, respectively (data not shown). The ORF method reduces the number of fragment pairs required to screen the search space by minimizing biologically irrelevant out-of-frame fragment pairings. From ORF filtering, 86% of the proteins were fully covered; however not all fragments were present in equal proportions. The fragment pairs were tested in two orientations since, theoretically, there is an equal chance for ligation with the activation domain (AD) µ-subunit of RNA polymerase or the DNA binding domain (DBD) λcI.

Complete coverage of the test space would result in 100% of amino acids in one protein being tested against 100% of amino acids in another. Here, the total possible test space between all covered proteins is shown in Fig. 2A and indicates that 47% of the possible test space was covered with 35% covered in both orientations. Six proteins were absent in the AD orientation (FOSL1, CSNK1E, CSNK2B, DKK1, DKK2, and CTNNBIP1), and two proteins were absent in the DBD orientation (DKK2 and JUN), meaning there were no fragments for those proteins in the specific orientation (Fig. 2A). Additionally, proteins containing an internal BstXI site are more likely to have poor or limited coverage. This restriction enzyme is required to ligate the fragment pairs into the pAVA plasmid [24]. The ORF filtering process (see methods) may also introduce a bias toward longer proteins and exacerbate poor coverage at the N- and C-termini [24]. Nevertheless, coverage of the search space yielded significant and interesting novel interactions.

**Figure 2.**
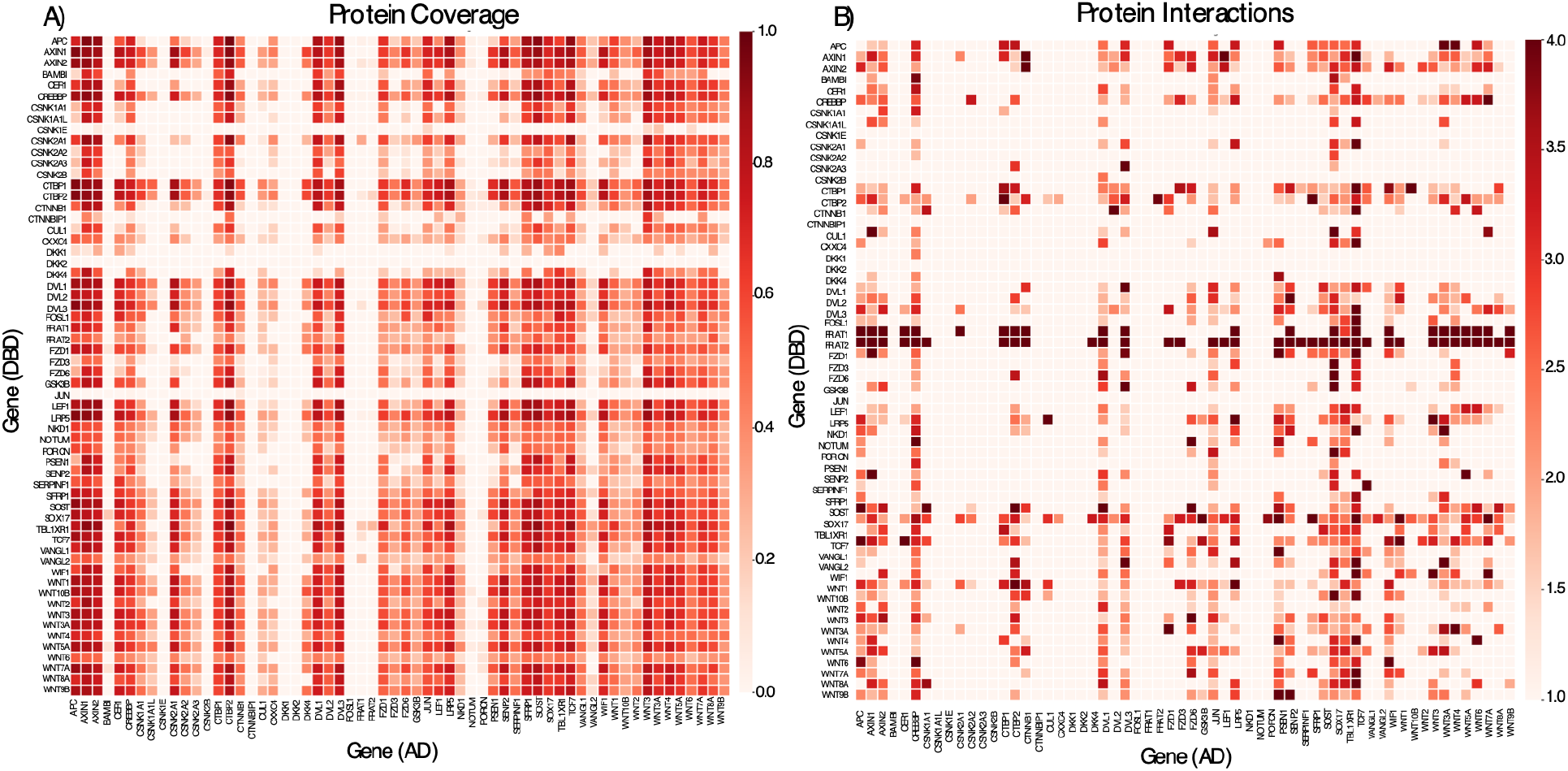
Sequence coverage and interaction heat map for Wnt pathway interactome. **A)** amino acid coverage of protein pairs. The 60 clones interrogated in this study are listed alphabetically. A value of 1 indicates 100% sequence coverage (red) for the protein pair, while a value of 0 indicates zero coverage for a given protein pair (light red). **B)** The protein pairs in panel A were tested for interactions by selection in the presence of 3-AT. Interactions were filtered based on Log_2_FC >1.5 and FDR <0.05. A value of 4 indicates a strong interaction with high Log_2_FC and minimum FDR, whereas a value of 1 indicates no interaction. Proteins paired with AD are represented on the x-axis, while proteins paired with DBD are represented on the y-axis.

### Significant pairs for protein-protein interactions

Fig. 2A shows a heat map indicating how well the protein-protein pairs were covered in this study. Interactions that pass significance filters, including FDR with multiple testing adjusted *p*-values, are shown in Fig. 2B. The color gradient in Fig. 2B represents the maximum detected Log_2_FC for paired fragments belonging to corresponding proteins. There were 597 PPIs identified in 2 mM 3-AT, 567 PPIs in 5 mM 3-AT growth conditions, and 461 PPIs in both 2 mM and 5 mM 3-AT. 106 PPIs were recovered exclusively in 5 mM 3-AT, and a majority (103 of 106) were present in 2 mM conditions near the cutoff in either FDR or Log_2_FC. These interactions present in the more stringent 5 mM but absent in 2 mM were borderline and deeper sequencing would likely recover them in 2 mM 3-AT conditions.

74 known interactions were recovered in this study (Table S2) [27]. Half of these were detected in both orientations (meaning AD-DBD and DBD-AD pairings) and have multiple unique fragment starting points present in 2 mM and 5 mM 3-AT conditions (Table S2). Fragments with multiple starting points (i.e., interacting fragments overlap) narrow the expected interaction region(s) between the proteins. Additionally, fragments interacting in AD-DBD and DBD-AD orientations increase the evidence that the interaction is real and not a false positive. Additional confidence is added for proteins that appear in two different libraries (unique transformation events) and interactions found in 2 mM and 5 mM 3-AT growth conditions.

### Detection of previously known APC and β-catenin complex

APC|AXIN1 and APC|CTNNB1 interactions were recovered in this study along with other well-established interactions, including GSK3β|AXIN1, GSK3β|AXIN2 [28], LRP5|GSK3β [29], LRP5|AXIN1 [30], LRP5|CTNNB1 [31], AXIN|CTNNB1 [28], CREBBP|CTNNB1 [32], and CTBP2|CTNNB1 [33] (Table S2). The APC|GSK3β interaction was not recovered (Fig. 2B), even though it was covered in both orientations (Fig. 2A).

### Detection of previously known interactions with nuclear transcription factors

In this study, known nuclear transcription factors interactions between CTNNB1|TCF7 [34], CTBP2|TCF7 [35], CTNNB1|SOX17 [18], JUN|CREBBP [34], JUN|CTBP2 [36], JUN|LRP5 [37], and FOSL1|CREBBP [36] (Table 1; Table S2) were recovered.

**Table 1.** List of nuclear protein interactions detected for APC and CTNNB1 using AVA-Seq. Columns 1 and 2 list Protein 1 and Protein 2, which are the tested protein pair. The following two columns, Orient 1 and Orient 2, show how many times the protein pair is detected in each orientation (Orient 1 is AD-associated; Orient 2 is DBD-associated) and then followed by a 3-AT competitive inhibitor condition to determine the number of pairs detected in 2 mM vs. 5 mM. Significant interactions filtered by Log_2_FCmax and FDRmin values. The number of libraries shows if the pairs are captured in a single library or both (with 2 being the maximum). The unique fragment pairs represent the number of unique fragments captured for each protein pair. The APID concludes if the pairs are novel (0) or known (1) previously from Agile Protein Interactomes DataServer (APID; [27]

| Protein1 | Protein2 | Orient1 | Orient2 | 2mM | 5mM | Log2FC<sub>max</sub> | FDR<sub>min</sub> | Library | Unique Pairs | APID |
| --- | --- | --- | --- | --- | --- | --- | --- | --- | --- | --- |
| AXIN1 | APC | 227 | 64 | 159 | 132 | 8.820432722 | 0 | 2 | 201 | 1 |
| APC | AXIN2 | 30 | 122 | 77 | 75 | 7.520242693 | 2.10E-97 | 2 | 123 | 1 |
| CTNNB1 | CTBP1 | 2 | 0 | 1 | 1 | 4.103357711 | 2.20E-25 | 1 | 1 | 1 |
| CTNNB1 | TBL1XR1 | 1 | 0 | 0 | 1 | 1.968622264 | 3.41E-23 | 1 | 1 | 1 |
| CTBP1 | APC | 73 | 5 | 46 | 32 | 6.198931965 | 1.52E-21 | 2 | 67 | 1 |
| AXIN2 | CTNNB1 | 9 | 2 | 9 | 2 | 6.050628355 | 6.77E-20 | 2 | 9 | 1 |
| CREBBP | APC | 280 | 79 | 147 | 212 | 5.577882812 | 3.70E-17 | 2 | 307 | 1 |
| TCF7 | APC | 34 | 17 | 22 | 29 | 5.891080664 | 1.45E-14 | 2 | 41 | 0 |
| DVL3 | APC | 127 | 25 | 62 | 90 | 5.171919508 | 5.19E-11 | 2 | 125 | 0 |
| CTNNB1 | CTBP2 | 5 | 14 | 8 | 11 | 3.257456516 | 1.33E-10 | 2 | 19 | 1 |
| DVL1 | APC | 50 | 10 | 31 | 29 | 4.154109344 | 1.12E-09 | 2 | 55 | 1 |
| CTNNB1 | AXIN1 | 7 | 6 | 8 | 5 | 5.025473084 | 3.20E-09 | 2 | 12 | 1 |
| APC | JUN | 25 | 0 | 10 | 15 | 5.5769084 | 5.80E-05 | 2 | 19 | 0 |
| SOX17 | APC | 99 | 25 | 49 | 75 | 5.453437899 | 5.04E-09 | 2 | 87 | 0 |
| DVL2 | APC | 30 | 0 | 15 | 15 | 5.102993509 | 2.17E-07 | 2 | 26 | 1 |
| CTNNB1 | CREBBP | 12 | 2 | 8 | 6 | 3.038695645 | 6.17E-07 | 2 | 14 | 1 |
| CTNNB1 | SOX17 | 7 | 8 | 5 | 10 | 5.184004171 | 1.26E-05 | 2 | 12 | 1 |
| FOSL1 | APC | 13 | 0 | 6 | 7 | 3.353875751 | 6.85E-05 | 2 | 8 | 0 |
| DVL1 | CTNNB1 | 3 | 3 | 5 | 1 | 3.155612862 | 0.00041615 | 2 | 5 | 1 |
| DVL3 | CTNNB1 | 4 | 4 | 5 | 3 | 3.966139708 | 0.00105864 | 2 | 8 | 1 |
| TCF7 | CTNNB1 | 1 | 2 | 0 | 3 | 2.067171582 | 0.00728363 | 1 | 3 | 1 |
| CTNNB1 | CSNK1A1 | 1 | 0 | 1 | 0 | 3.500906308 | 0.00888600 | 1 | 1 | 1 |
| CTNNB1 | DVL2 | 2 | 0 | 1 | 1 | 4.682673739 | 0.00891623 | 1 | 1 | 1 |

### Novel interactions of APC with nuclear transcription factors

APC interacted with several nuclear proteins, of which six interactions are known and four are novel (Table 1). The novel binding partners for APC and their localized interaction regions identified in this study (Table 1) include JUN (Fig 3A), FOSL1 (Fig 3B), SOX17 (Fig 3C), and TCF7 (Fig 3D). Our data show APC binds to JUN, FOSL1, and TCF7 in the same interaction region required for CTNNB1 binding (Fig. S6). Other known nuclear protein interactions are detected for both APC and CTNNB1 (Table 1) [38, 17, 37, 39].

**Figure 3.**
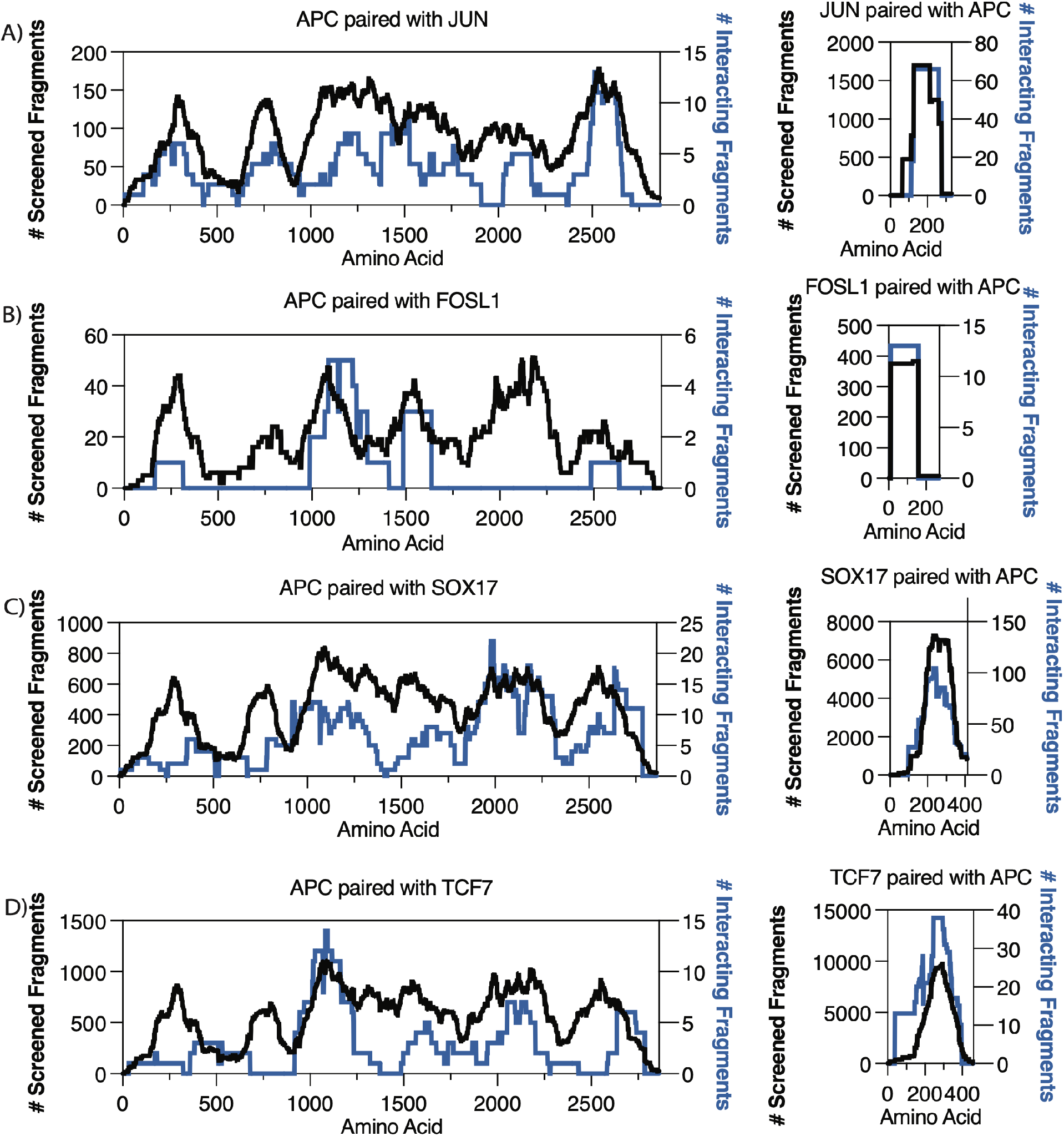
High-resolution interaction mapping of APC with transcription factors. The y-axis represents the total screened fragments (left; black trace) and the number of interacting fragments (right; blue trace). The x-axis represents protein length by amino acids. **A)** APC interaction region detected with JUN protein. **B)** APC interaction region detected with FOSL1 protein. **C)** APC interaction detected with SOX17 protein. **D)** APC interaction detected with TCF7 protein.

The fragment interactions between APC|JUN are present only in one orientation since JUN is only fused with the AD (Fig. 2A), with 19 unique interacting fragments (Table 1). The fragment interactions between APC|FOSL1 are present only in one orientation since FOSL1 is only fused with the DBD (Fig. 2A), with eight unique interacting fragments (Table 1). For APC|SOX17, 87 unique interacting fragment pairs were recovered in both orientations (Table 1). For APC|TCF7, 41 unique interacting fragment pairs were recovered in both orientations (Table 1).

### Secondary validation of APC interactions with nuclear transcription factors

APC fragments used in the pull-down assays were designed to cover the most prevalent interaction regions observed between APC and the transcription factors (Fig 4). In total, four APC fragments were tested against full-length and fragments of TCF7, JUN, FOSL1, and SOX17 proteins.

**Figure 4.**
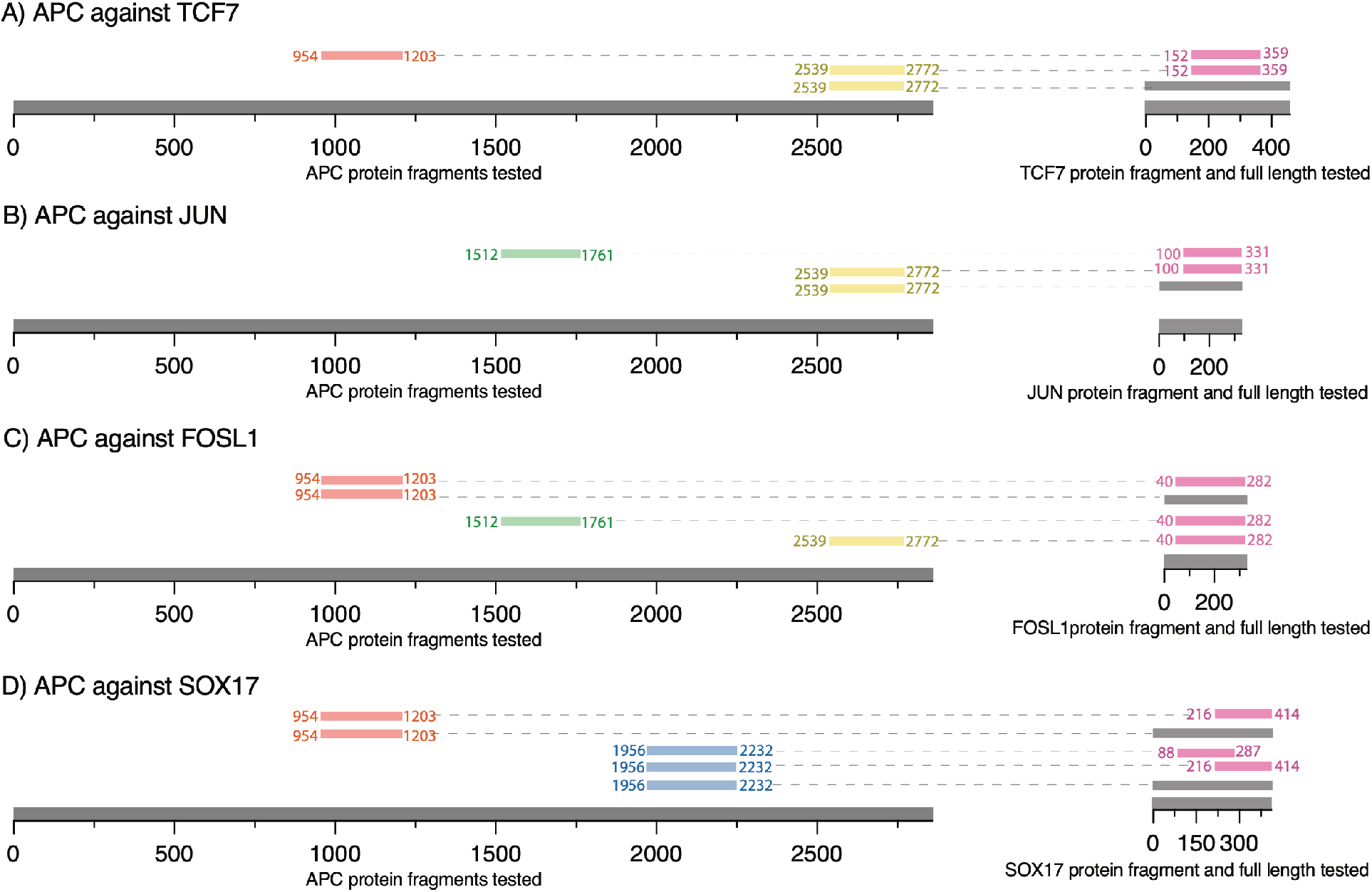
Secondary validation using pull-down of protein pairs. The exact fragment residues are indicated with superscript, with full-length represented by superscript ‘FL’. The first fragment, APC^954-1203^ (red), spanned the 15R region; the second fragment, APC^1512-1761^ (green), spanned the fourth 20R region and SAMP1-2 regions; the third fragment, APC^1956-2232^ (blue), covered the sixth and seventh 20R region and the SAMP3 repeat; the fourth fragment APC^2539-2772^ (yellow) covered the EB1 domain. **A)** APC^954-1203^ (red) was tested against TCF7^152-359^ (pink). APC^2539-2772^ (yellow) was tested against TCF7^152-359^ and TCF7^FL^ (grey). **B)** APC^1512-1761^ (green) was tested against JUN^100-331^ (pink). APC^2539-2772^ (yellow) was tested against JUN^100-331^ (pink) and JUN^FL^ (grey). **C)** APC^954-1203^ (red) was tested against FOSL1^40-282^ (pink) and FOSL1^FL^ (grey). APC^1512-1761^ (green) was tested against FOSL1^40-282^ (pink). APC^2539-2772^ (yellow) was tested against FOSL1^40-282^ (pink) as a negative control. **D)** APC^954-1203^ (red) was tested against SOX17^216-414^ (pink) and SOX17^FL^ (grey). APC^1956-2232^ (blue) was tested against SOX17^88-287^, SOX17^216-414^ (pink), and SOX17^FL^.

The negative control used for all pull-down experiments was MBP without a fusion protein (Fig. 5 lane 4; Fig. 6 lane 6; Fig. S3 lane 1; Fig. S4 lane 4; Fig. S5 lane 3) to ensure the GST-APC fragments did not interact with MBP itself. Based on our interaction results (Fig. 3D) and pull-down experiments (Fig. 5 lane 3; Fig. 6 lanes 4-5), APC protein directly interacts with TCF7. MBP-TCF7^152-359^ pulled down GST-APC^954- 1203^ (Fig. 5 lane 3) and GST-APC^2539-2772^ (Fig. 6 lane 4), indicating a direct interaction of APC with TCF7. MBP-TCF7^152-359^ and MBP-TCF7^FL^ pulled down the GST-APC^2539-2772^ fragment (Fig. 6 lane 4-5). The interaction of GST-APC^2539-2772^ had a similar signal intensity to MBP-TCF7^152-359^ and MBP-TCF7^FL^ (Fig. 6 lanes 4-5).

**Figure 5.**
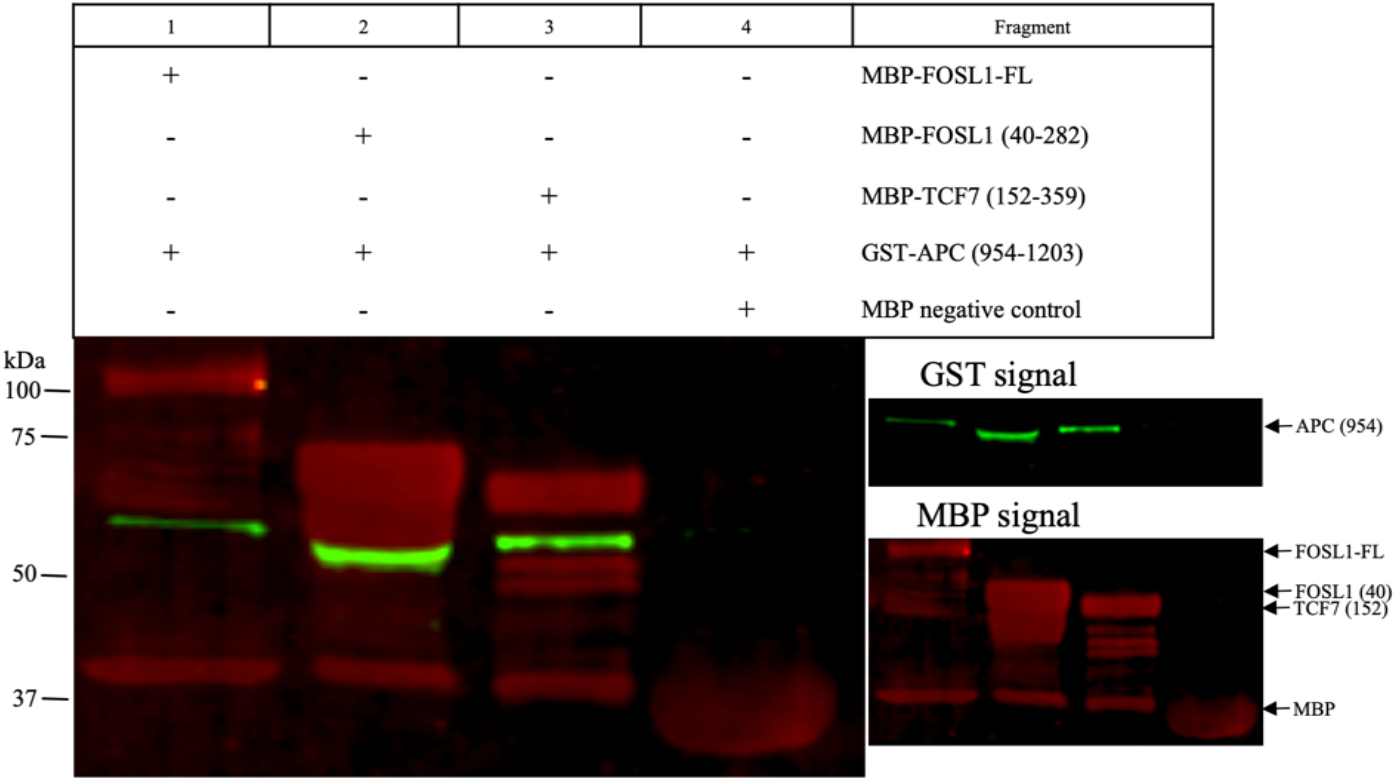
Pull-down of APC^954-1203^ containing the 15R region. GST-APC^954-1203^ was tested against the following MBP tagged proteins: Lane 1: FOSL1^FL^, Lane 2: FOSL1^40-282^, Lane 3: TCF7^152-359^, and Lane 4: MBP alone (negative control). All proteins expressed with MBP show a leaky expression of MBP as indicated by a red band of 40.3 kDa and present in protein expression gels (Fig. S1BD). The signal of MBP alone in all samples (lanes 1-3) represents the binding of MBP protein to the amylose resin. Samples FOSL1^FL^, FOSL1^40-282^, and TCF7^152-359^ show fragmentation represented in the gel by multiple bands below the expected target protein. Arrow points to the expected molecular weight of the target protein/fragment (green signal GST; red signal MBP). Each pull-down experiment was conducted in triplicate.

**Figure 6.**
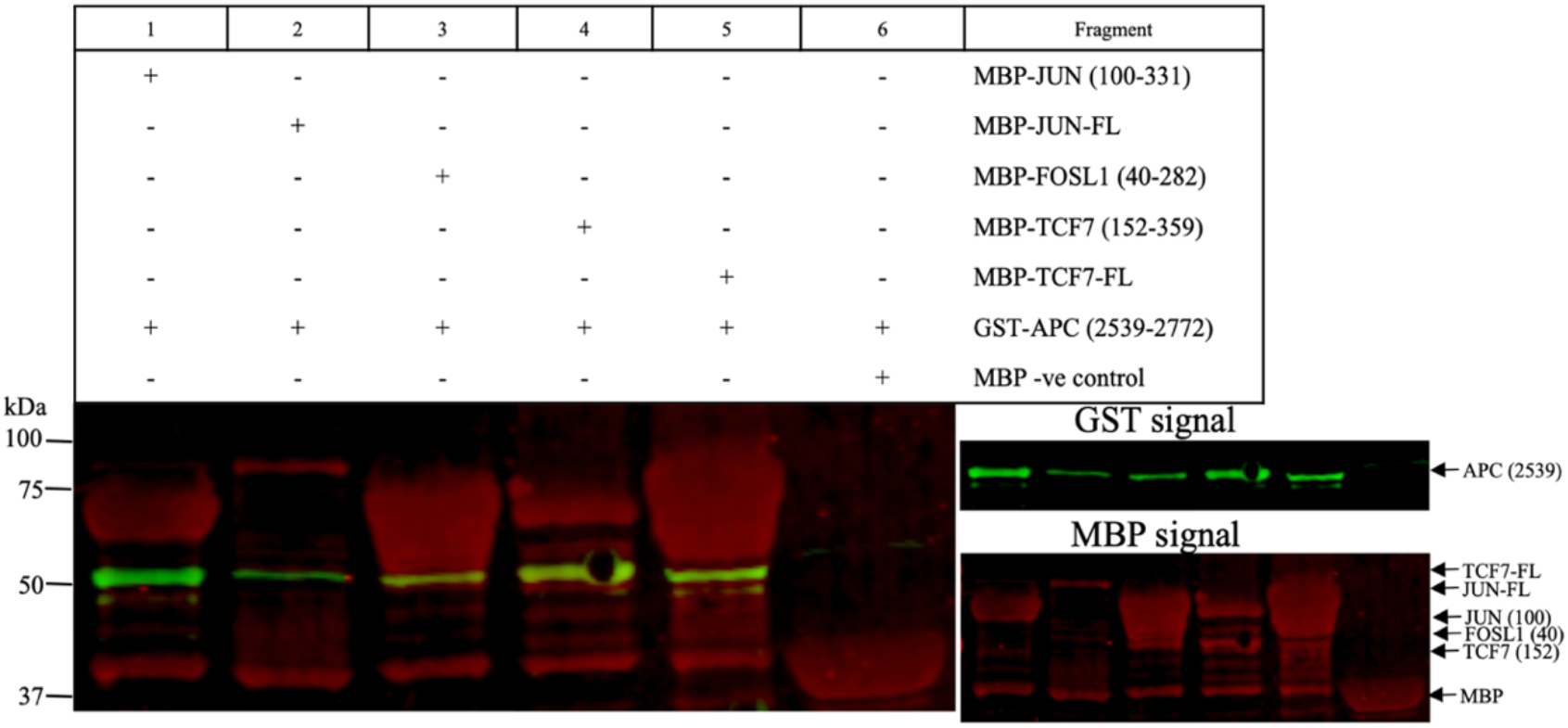
Pull-down of GST-APC^2539-2772^ containing the EB1 domain. GST-APC^2539-2772^ was tested against the following MBP tagged proteins: JUN^100-331^ (lane 1), JUN^FL^ (lane 2), FOSL1^40-282^ (lane 3), TCF7^152-359^ (lane 4), TCF7^FL^ (lane 5), and MBP alone (negative control; lane 6). The signal of MBP alone in all the samples (lanes 1-5) represents the binding of cleaved MBP protein to the amylose resin, likely resulting from leaky expression or cleavage during purification. The TCF7^152-359^ and TCF7^FL^ constructs show fragmentation represented by multiple bands below the expected target protein. An arrow indicates the expected molecular weight of the target protein/fragment (green signal GST; red signal MBP).

APC directly interacts with FOSL1 (Fig. 5 lane 1-2; Fig. 6 lane 3; Fig. S3 lane 3). The interaction of MBP-FOSL1^FL^ with GST-APC^954-1203^ (Fig. 5 lane 1) is weaker compared to the fragment MBP-FOSL1^40-282^ interaction (Fig. 5 lane 2) based on the signal intensity obtained with GST-APC^954-1203^. The MBP-FOSL1^40- 282^ pulled down GST-APC^2539-2772^ (Fig. 6 lane 3). Also, APC interacted directly with JUN (Fig. 6 lane 1-2: Fig. S3 lane 2). The interaction of MBP-JUN^FL^ with GST-APC^2539-2772^ appears weaker than the MBP-JUN^100-331^ interaction (Fig. 6 lane 2). The negative control (MBP without a fusion protein) could not pull down GST-APC^2539-2772^.

MBP-JUN^100-331^ and MBP-FOSL1^40-282^ pulled down GST-APC^1512-1761^ (Fig. S3 lanes 2-3, respectively), suggesting these transcription factors bind this region of APC. Pull-down validation for SOX17 utilized MBP-SOX17^88-287^, MBP-SOX17^216-414^, and MBP-SOX17^FL^ to pull down GST-APC^1956-2232^ (Fig. S4 lanes 1-3, respectively). MBP-SOX17^FL^ and MBP-SOX17^216-414^ proteins pulled down GST-APC^954-1203^ (Fig. S5 lanes 1-2, respectively). The interaction of MBP-SOX17^216-414^ appears to bind GST-APC^1956-2232^ weaker than MBP-SOX17^88-287^ and MBP-SOX17^FL^, which is indicated by the GST signal (Fig. S4 lane 2).

## Discussion

APC is an integral protein of the Wnt β-catenin signaling pathway and forms a complex with several proteins in the cytosol, including AXIN1, CTNNB1, and GSK3β. For the first time, we applied the AVA-Seq method to determine protein-protein interactions in the Wnt pathway. Adding another dimension to previously observed localization of APC in the nucleus, we found APC interacted with JUN, FOSL1, TCF7, and SOX17 transcription factors (Table 1). Additionally, our data indicate enrichment of interacting fragments in the known interaction region for APC|AXIN1 [40], APC|CTNNB1 [40], AXIN1|CTNNB1 [38], and CTNNB1|JUN [8]. The APC|GSK3β [41] interaction was not recovered here even though fragment pairs covered the proteins. This is not surprising as phosphorylation is required for the interaction to occur, and post-translational modification does not happen in the bacterial system. It was interesting to investigate APC interactions with nuclear transcription factors which could expand on the functional importance and role of APC in cancer (Fig. 3; Table 1). We validated novel APC interactions with the transcription factors using a secondary assay (Fig. 4).

A previous study showed that APC competes with TCF7 for CTNNB1 binding [17] (Fig. 7 pathway B). However, the role of direct binding of APC with TCF7 is not yet well understood. APC is known to form a complex with Src associated with mitosis (Sam68) protein to regulate alternative splicing of TCF1 which is also known as TCF7 [48]. Upon LOF of APC, the splice variant TCF1E accumulates and induces expression of the Wnt target gene [48]. Moreover, TCF7 mRNA expression positively correlated with the mutation of CTNNB1, whereas APC levels were unaffected [48]. Several reports have shown both TCF7 and CTNNB1 are responsible for tumor formation and metastasis [47, 48]. Despite the correlation between CTNNB1 mutation with TCF7 accumulation, the role of APC as a possible mediator has not been investigated.

**Figure 7.**
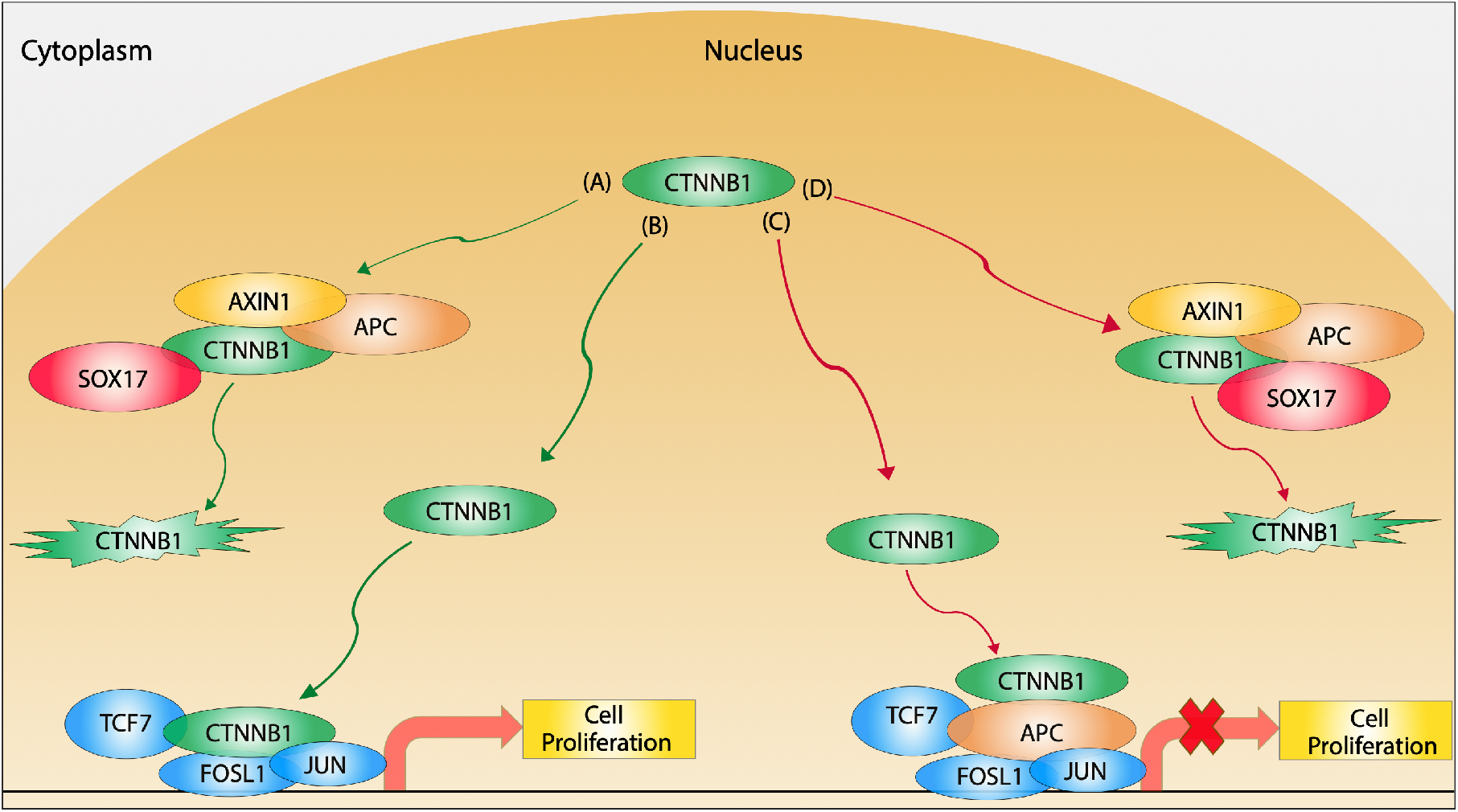
Wnt pathway APC/CTNNB1 nuclear model known and proposed. Known nuclear model: **A)** For nuclear CTNNB1 degradation, both APC anchored with AXIN [45] binds to CTNNB1 along with SOX17 [18] and marks it for ubiquitin degradation. **B)** When CTNNB1 enters the nucleus, it could proceed and bind transcription factors TCF7 [46], JUN, and FOSL1 [8] to initiate cell proliferation. Proposed nuclear model: **C)** When CTNNB1 enters the nucleus, it could be inhibited through APC binding to TCF7, FOSL1, and JUN transcription factors. **D)** For nuclear CTNNB1 degradation, APC, AXIN1, and SOX17 bind to CTNNB1. However, this might require the binding of SOX17 to both CTNNB1 (known) and APC (novel) to mark CTNNB1 for ubiquitin degradation. Green pathway (known); red pathway (a proposed mechanism).

APC interacted with TCF7 in the region required for CTNNB1 binding (Fig. S6D) through multiple unique fragments indicating the robustness of the interaction in our system. This interaction region was further validated with pull-down assays (Fig. 3D; Fig. 5 lane 3). In addition, the interaction of TCF7 with 15R and 20R repeats of APC might serve a similar function to APC’s interaction with CTNNB1 as it’s known APC binds CTNNB1 tightly through the 15R repeat [49] and requires the 20R repeat to down-regulate CTNNB1 [49]. The 15R region of APC is critical for C-terminal binding protein (CTBP) to down-regulate TCF [23] and most CRC is associated with mutations in the 20R repeat [42]. We propose APC binds TCF7 to repress and down-regulate the CTNNB1|TCF7 interaction and not just compete for CTNNB1 binding alone (Fig. 7 pathway C).

CTNNB1 drives proliferation through direct binding to FOSL1 and JUN [8, 49]. APC LOF is correlated with JUN over-expression [50]. A recent study showed that fat-1 transgenic mice had a down-regulated expression of APC and FOSL1 compared to wild-type [51]. Even with these gene expression correlations, there has not been a report of direct binding of APC with the two proteins. To the best of our knowledge, this is the first report of APC interacting with FOSL1 and JUN through both two-hybrid and pull-down assays. To our surprise, APC interacted with FOSL1 and JUN in the region required for CTNNB1 binding (Fig. S6AB domain: white box). Based on our findings, we suggest APC could inhibit CTNNB1 gene expression by directly binding JUN and FOSL1 transcription factors (Fig. 7 pathway C) since APC binds to both transcription factors in the same region required for CTNNB1 binding.

It has been reported that SOX17 is a vital target in CRC since it targets both nuclear TCF7 and CTNNB1 for degradation [5, 17, 52, 53, 54] (Fig. 7 pathway A). SOX17 functions similarly to APC by acting as a tumor suppressor and negatively regulates the Wnt pathway [52]. Further, more than 80% of cancer patients have methylation of SOX17 promoter, which is negatively associated with the accumulation of nuclear CTNNB1 [53, 54]. Aside from detecting the novel SOX17|APC interaction, the known interactions of SOX17|CTNNB1 and SOX17|TCF7 [18] were recovered (Table 1). In our findings, APC is mostly bound to the central region of SOX17 and not in the region of CTNNB1 contact (Fig. S6C). Furthermore, since APC and SOX17 are tumor suppressor genes, we conclude that APC’s interaction with SOX17 might enhance CTNNB1 degradation in the nucleus (Fig. 7 pathway D).

Here we have shown that APC interacts with nuclear transcription factors JUN, FOSL1, TCF7, and SOX17 in the bacterial two-hybrid AVA-Seq method and validation pull-down assays using both truncated and full-length proteins. We suggest a possible mechanism of nuclear APC activity to bind TCF7, JUN, and FOSL1 in the region required for CNTNB1 binding, while nuclear APC binds SOX17 to enhance CTNNB1 degradation. This information is an important addition to previous observations of APC localizing to the nucleus and helps shed light on APC nuclear function. These interactions may offer new drug targets to reduce tumor formation and malignancy. We plan in the future to focus on understanding how mutations in the identified contact regions might affect the protein interactions.

## Acknowledgments and funding

This research was supported by funding from Qatar Foundation to Weill Cornell Medicine in Qatar in the form of the BMRP2 grant.

## Ethical approval statement

No humans or animals were used for this study; thus, no ethical approval or patient consent is required.

## Data sharing and data accessibility

Sequences were deposited to the Sequence Read Archive of NCBI under the BioProject ID PRJNA841056.

## Conflict of Interest

The authors declare there are no conflicts or competing interests.

## Author Contributions

N.M.A. and J.A.M. conceived the idea and designed the study; N.M.A., J.A.M., and Y.A.M. collected the data; N.M.A., J.A.M., J.A., and S.S.-R. analyzed the data; N.M.A., J.A.M., J.A., and S.S.-R. wrote the manuscript.

## Supplementary material

**Figure S1**. Expression gels: TCF7, JUN, FOSL1 protein with MBP, and APC protein with GST tag.

**Figure S2**. Expression gel: SOX17 protein with MBP tag and MBP tag negative control.

**Figure S3**. Pull-down GST-APC^1512-1761^ with MBP-JUN, MBP-FOSL1, and MBP negative control.

**Figure S4**. Pull-down GST-APC^1956-2232^ with MBP-SOX17 and MBP negative control.

**Figure S5**. Pull-down GST-APC^954-1203^ with MBP-SOX17 and MBP negative control.

**Figure S6**. Interaction mapping showing domains for (APC, JUN, FOSL1, TCF7, SOX17) and CTNNB1 binding region on transcription factors.

**Table S1**. The 60 Wnt pathway clones from Genscript.

**Table S2**. Stringent known protein-protein interactions detected.

**Table S3**. List of tested fragments for pull-down.

## Notes

https://www.ncbi.nlm.nih.gov/bioproject/?term=PRJNA841056.

### Competing Interest Statement

The authors have declared no competing interest.

## References

1. Siegel, R.L., et al., Cancer statistics, 2022. CA Cancer J Clin, 2022. 72(1): p. 7–33.

2. Anderson, C.B., K.L. Neufeld, and R.L. White, Subcellular distribution of Wnt pathway proteins in normal and neoplastic colon. Proc Natl Acad Sci U S A, 2002. 99(13): p. 8683–8.

3. Neufeld, K.L. and R.L. White, Nuclear and cytoplasmic localizations of the adenomatous polyposis coli protein. Proc Natl Acad Sci U S A, 1997. 94(7): p. 3034–9.

4. Katoh, M., Canonical and non-canonical WNT signaling in cancer stem cells and their niches: Cellular heterogeneity, omics reprogramming, targeted therapy and tumor plasticity (Review). Int J Oncol, 2017. 51(5): p. 1357–1369.

5. Jeong, W.J., E.J. Ro, and K.Y. Choi, Interaction between Wnt/beta-catenin and RAS-ERK pathways and an anti-cancer strategy via degradations of beta-catenin and RAS by targeting the Wnt/beta-catenin pathway. NPJ Precis Oncol, 2018. 2(1): p. 5.

6. Guo, Y., et al., EHMT2 promotes the pathogenesis of hepatocellular carcinoma by epigenetically silencing APC expression. Cell Biosci, 2021. 11(1): p. 152.

7. Ota, R., et al., Integrated genetic and epigenetic analysis of cancer-related genes in non-ampullary duodenal adenomas and intramucosal adenocarcinomas. J Pathol, 2020. 252(3): p. 330–342.

8. Toualbi, K., et al., Physical and functional cooperation between AP-1 and beta-catenin for the regulation of TCF-dependent genes. Oncogene, 2007. 26(24): p. 3492–502.

9. Eferl, R. and E.F. Wagner, AP-1: a double-edged sword in tumorigenesis. Nat Rev Cancer, 2003. 3(11): p. 859–68.

10. Huang, T.S., S.C. Lee, and J.K. Lin, Suppression of c-Jun/AP-1 activation by an inhibitor of tumor promotion in mouse fibroblast cells. Proc Natl Acad Sci U S A, 1991. 88(12): p. 5292–6.

11. Jochum, W., et al., Increased bone formation and osteosclerosis in mice overexpressing the transcription factor Fra-1. Nat Med, 2000. 6(9): p. 980–4.

12. Young, M.R., et al., Transgenic mice demonstrate AP-1 (activator protein-1) transactivation is required for tumor promotion. Proc Natl Acad Sci U S A, 1999. 96(17): p. 9827–32.

13. Korinek, V., et al., Constitutive transcriptional activation by a beta-catenin-Tcf complex in APC-/-colon carcinoma. Science, 1997. 275(5307): p. 1784–7.

14. Hrckulak, D., et al., TCF/LEF Transcription Factors: An Update from the Internet Resources. Cancers (Basel), 2016. 8(7).

15. Tang, W., et al., A genome-wide RNAi screen for Wnt/beta-catenin pathway components identifies unexpected roles for TCF transcription factors in cancer. Proc Natl Acad Sci U S A, 2008. 105(28): p. 9697–702.

16. Nateri, A.S., B. Spencer-Dene, and A. Behrens, Interaction of phosphorylated c-Jun with TCF4 regulates intestinal cancer development. Nature, 2005. 437(7056): p. 281–5.

17. Sinner, D., et al., Sox17 and Sox4 differentially regulate beta-catenin/T-cell factor activity and proliferation of colon carcinoma cells. Mol Cell Biol, 2007. 27(22): p. 7802–15.

18. Zorn, A.M., et al., Regulation of Wnt signaling by Sox proteins: XSox17 alpha/beta and XSox3 physically interact with beta-catenin. Mol Cell, 1999. 4(4): p. 487–98.

19. Choi, S.G., et al., Maximizing binary interactome mapping with a minimal number of assays. Nat Commun, 2019. 10(1): p. 3907.

20. Schaefer-Ramadan, S., et al., Novel protein contact points among TP53 and minichromosome maintenance complex proteins 2, 3, and 5. Cancer Med, 2022.

21. Dove, S.L. and A. Hochschild, A bacterial two-hybrid system based on transcription activation. Methods Mol Biol, 2004. 261: p. 231–46.

22. Andrews, S.S., et al., High-resolution protein-protein interaction mapping using all-versus-all sequencing (AVA-Seq). J Biol Chem, 2019. 294(30): p. 11549–11558.

23. Hamada, F. and M. Bienz, The APC tumor suppressor binds to C-terminal binding protein to divert nuclear beta-catenin from TCF. Dev Cell, 2004. 7(5): p. 677–85.

24. Schaefer-Ramadan, S., et al., Scaling-up a fragment-based protein-protein interaction method using a human reference interaction set. Proteins, 2022. 90(4): p. 959–972.

25. Buchfink, B., C. Xie, and D.H. Huson, Fast and sensitive protein alignment using DIAMOND. Nat Methods, 2015. 12(1): p. 59–60.

26. Hebditch, M., et al., Protein-Sol: a web tool for predicting protein solubility from sequence. Bioinformatics, 2017. 33(19): p. 3098–3100.

27. Alonso-Lopez, D., et al., APID interactomes: providing proteome-based interactomes with controlled quality for multiple species and derived networks. Nucleic Acids Res, 2016. 44(W1): p. W529–35.

28. Ikeda, S., et al., Axin, a negative regulator of the Wnt signaling pathway, forms a complex with GSK-3beta and beta-catenin and promotes GSK-3beta-dependent phosphorylation of beta-catenin. EMBO J, 1998. 17(5): p. 1371–84.

29. Mi, K., P.J. Dolan, and G.V. Johnson, The low density lipoprotein receptor-related protein 6 interacts with glycogen synthase kinase 3 and attenuates activity. J Biol Chem, 2006. 281(8): p. 4787–94.

30. Mao, J., et al., Low-density lipoprotein receptor-related protein-5 binds to Axin and regulates the canonical Wnt signaling pathway. Mol Cell, 2001. 7(4): p. 801–9.

31. Aripaka, K., et al., TRAF6 function as a novel co-regulator of Wnt3a target genes in prostate cancer. EBioMedicine, 2019. 45: p. 192–207.

32. Yu, W., et al., beta-Catenin Cooperates with CREB Binding Protein to Promote the Growth of Tumor Cells. Cell Physiol Biochem, 2017. 44(2): p. 467–478.

33. Kim, T.W., et al., Ctbp2-mediated beta-catenin regulation is required for exit from pluripotency. Exp Mol Med, 2017. 49(10): p. e385.

34. Benbrook, D.M. and N.C. Jones, Heterodimer formation between CREB and JUN proteins. Oncogene, 1990. 5(3): p. 295–302.

35. Zhang, W., et al., The intracellular NADH level regulates atrophic nonunion pathogenesis through the CtBP2-p300-Runx2 transcriptional complex. Int J Biol Sci, 2018. 14(14): p. 2023–2036.

36. Li, X., et al., Proteomic analyses reveal distinct chromatin-associated and soluble transcription factor complexes. Mol Syst Biol, 2015. 11(1): p. 775.

37. Larabee, J.L., et al., Adenomatous polyposis coli protein associates with C/EBP beta and increases Bacillus anthracis edema toxin-stimulated gene expression in macrophages. J Biol Chem, 2011. 286(22): p. 19364–72.

38. Xing, Y., et al., Crystal structure of a beta-catenin/axin complex suggests a mechanism for the beta-catenin destruction complex. Genes Dev, 2003. 17(22): p. 2753–64.

39. Matsumoto, S., et al., Binding of APC and dishevelled mediates Wnt5a-regulated focal adhesion dynamics in migrating cells. EMBO J, 2010. 29(7): p. 1192–204.

40. Eklof Spink, K., S.G. Fridman, and W.I. Weis, Molecular mechanisms of beta-catenin recognition by adenomatous polyposis coli revealed by the structure of an APC-beta-catenin complex. EMBO J, 2001. 20(22): p. 6203–12.

41. Rubinfeld, B., et al., Binding of GSK3beta to the APC-beta-catenin complex and regulation of complex assembly. Science, 1996. 272(5264): p. 1023–6.

42. Liu, J., et al., The third 20 amino acid repeat is the tightest binding site of APC for beta-catenin. J Mol Biol, 2006. 360(1): p. 133–44.

43. Hankey, W., W.L. Frankel, and J. Groden, Functions of the APC tumor suppressor protein dependent and independent of canonical WNT signaling: implications for therapeutic targeting. Cancer Metastasis Rev, 2018. 37(1): p. 159–172.

44. Barth, A.I., K.A. Siemers, and W.J. Nelson, Dissecting interactions between EB1, microtubules and APC in cortical clusters at the plasma membrane. J Cell Sci, 2002. 115(Pt 8): p. 1583–90.

45. Rosin-Arbesfeld, R., F. Townsley, and M. Bienz, The APC tumour suppressor has a nuclear export function. Nature, 2000. 406(6799): p. 1009–12.

46. Molenaar, M., et al., XTcf-3 transcription factor mediates beta-catenin-induced axis formation in Xenopus embryos. Cell, 1996. 86(3): p. 391–9.

47. Atay, S., Integrated transcriptome meta-analysis of pancreatic ductal adenocarcinoma and matched adjacent pancreatic tissues. PeerJ, 2020. 8: p. e10141.

48. Juca, C.E.B., et al., Impact of the Canonical Wnt Pathway Activation on the Pathogenesis and Prognosis of Adamantinomatous Craniopharyngiomas. Horm Metab Res, 2018. 50(7): p. 575–581.

49. Rubinfeld, B., et al., Loss of beta-catenin regulation by the APC tumor suppressor protein correlates with loss of structure due to common somatic mutations of the gene. Cancer Res, 1997. 57(20): p. 4624–30.

50. Saadeddin, A., et al., The links between transcription, beta-catenin/JNK signaling, and carcinogenesis. Mol Cancer Res, 2009. 7(8): p. 1189–96.

51. Warner, D.R., et al., Beneficial effects of an endogenous enrichment in n3-PUFAs on Wnt signaling are associated with attenuation of alcohol-mediated liver disease in mice. FASEB J, 2021. 35(2): p. e21377.

52. Jia, Y., et al., SOX17 antagonizes WNT/beta-catenin signaling pathway in hepatocellular carcinoma. Epigenetics, 2010. 5(8): p. 743–9.

53. Wang, L., et al., SOX17 Antagonizes the WNT Signaling Pathway and is Epigenetically Inactivated in Clear-Cell Renal Cell Carcinoma. Onco Targets Ther, 2021. 14: p. 3383–3394.

54. Zhou, W., et al., SOX17 Inhibits Tumor Metastasis Via Wnt Signaling In Endometrial Cancer. Onco Targets Ther, 2019. 12: p. 8275–8286.

