## Supplemental Information for "Identifying novel interactions of the colon-cancer related APC protein with Wnt-pathway nuclear transcription factors"

### Supplemental Figures:

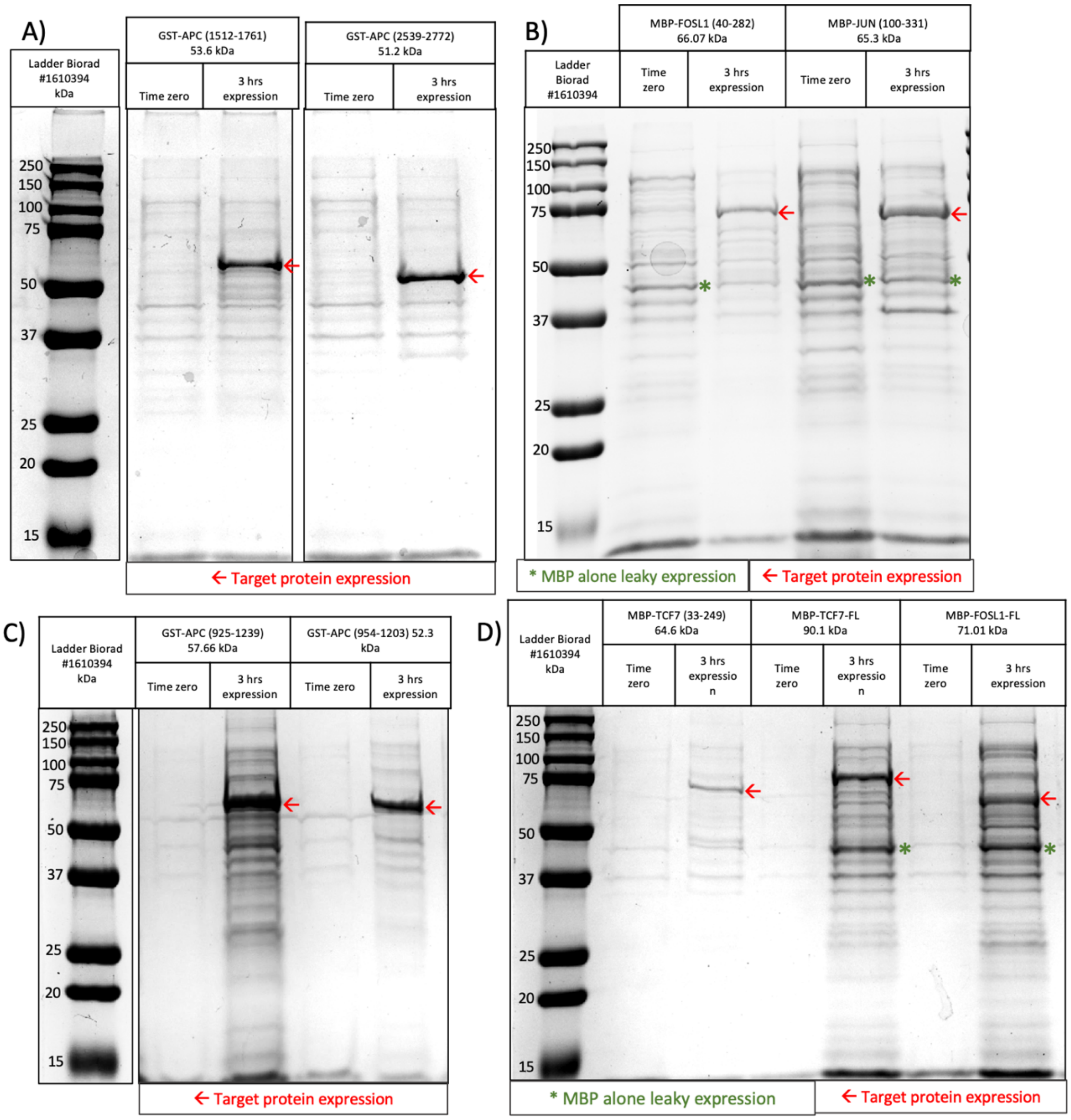


**Supplemental Figure 1: Protein expression gels of full-length and fragments used for pull-down experiments. A)** two fragments of APC were expressed with GST tag for 3 hours GST-APC$^{1512\text{-}1761}$ and GST-APC$^{2539\text{-}2772}$. Time zero indicates the start point for protein expression cells OD = 0.5-0.6, then followed by 3 hours of expression with IPTG. **B)** MBP-FOSL1$^{40\text{-}282}$ and MBP-JUN$^{100\text{-}331}$ at time zero, followed by 3 hours of expression with IPTG. **C)** GST-APC$^{938\text{-}1239}$ and GST-APC$^{954\text{-}1203}$ at times zero followed by 3 hours expression with IPTG. **D)** MBP-TCF7$^{152\text{-}359}$, MBP-TCF7$^{FL}$, and MBP-FOSL1$^{FL}$ expression at time zero, followed by 3 hours expression with IPTG. The red arrow (←) points to target protein fragment expression, while (*) is the leaky expression of MBP seen in B, and D. Protein loading concentration 40-60 µg.

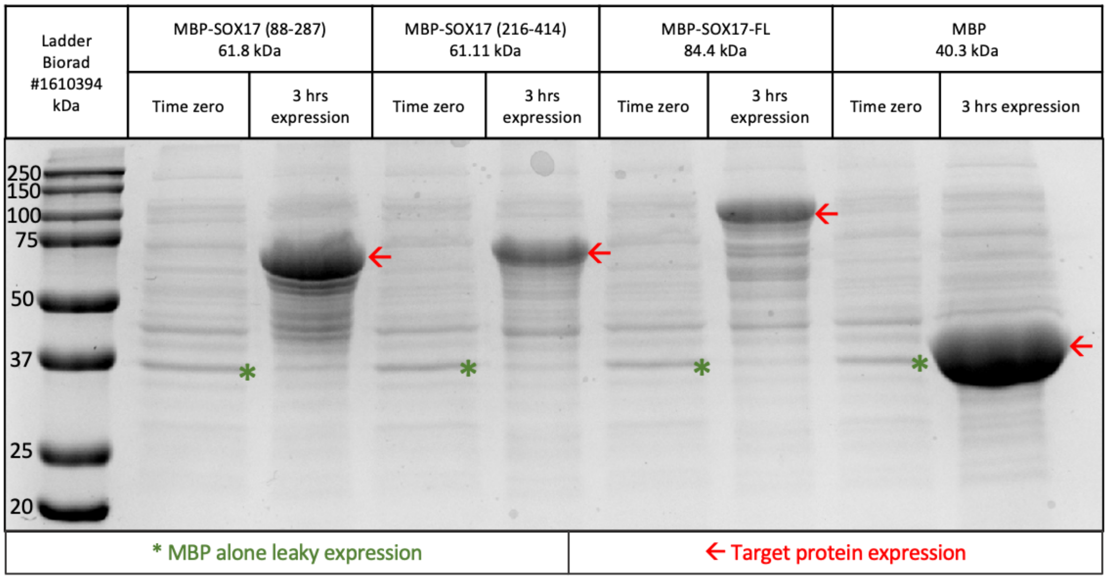


**Supplemental Figure 2: Protein expression of SOX17 pull-down.** Three fragments of SOX17 expressed with MBP tag for 3 hours SOX17$^{88\text{-}247}$; SOX17$^{216\text{-}415}$; SOX17$^{FL}$; along with MBP tag alone (negative control for pull-down). Time zero indicates the start point for protein expression cells OD = 0.5-0.6, then followed by 3 hours expression with IPTG. The red arrow (←) points to target protein fragment expression, while (*) is the leaky expression of MBP. Protein loading concentration 40-60 μg.

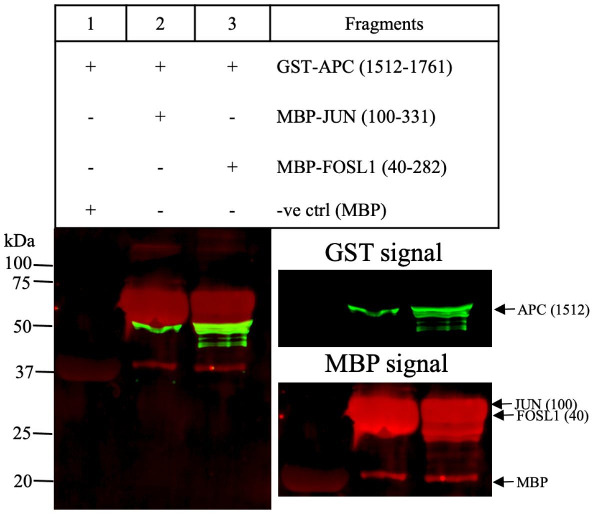


**Supplemental Figure 3: Pull-down of GST-APC$^{1512\text{-}1761}$ containing the fourth 20R region, SAMP1, and SAMP2 repeats.** GST-APC$^{1512\text{-}1761}$ was tested against the following MBP tagged proteins: Lane 1: MBP alone (negative control), Lane 2: JUN$^{100\text{-}331}$, Lane 3: FOSL1$^{40\text{-}282}$. All proteins expressed with MBP show a leaky expression of MBP as indicated by a red band 40.3 kDa and present in protein expression gels (Fig. S1BD). The signal of MBP alone in the samples (well 2-3) represents the binding of MBP protein to the amylose resin. Lane 3: sample GST-APC$^{1512\text{-}1761}$ shows fragmentation represented in the gel by multiple bands below the expected target protein. Arrows point to the expected molecular weight of the target protein fragment (green signal GST; red signal MBP).

| 1 | 2 | 3 | 4 | Fragment |
|---|---|---|---|---|
| + | - | - | - | MBP-SOX17 (88-287) |
| - | + | - | - | MBP-SOX17 (216-415) |
| - | - | + | - | MB-SOX17-FL |
| + | + | + | + | GST-APC (1956-2232 ) |
| - | - | - | + | MBP negative control |

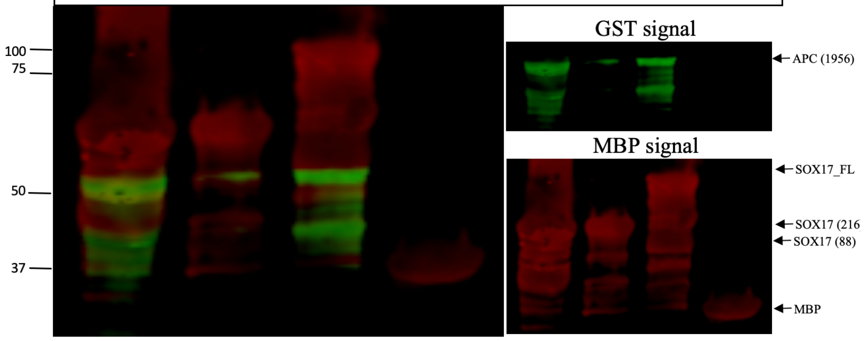


**Supplemental Figure 4: Pull-down of the GST-APC$^{1956-2232}$ which contain the sixth 20R region and SAMP3 repeat.** GST-APC$^{1956-2232}$ was tested against the following MBP tagged proteins: Lane 1: SOX17$^{88-287}$, Lane 2: SOX17$^{216-415}$, Lane 3: SOX17$^{FL}$, and Lane 4: MBP alone (negative control). Lane 1 and 3: GST-APC$^{1956-2232}$ shows fragmentation represented in the gel by multiple bands below the expected target protein. The arrows point to the expected molecular weight of the target protein fragment (green signal GST; red signal MBP).

| 1 | 2 | 3 | Fragment |
|---|---|---|---|
| + | - | - | MBP-SOX17-FL |
| - | + | - | MBP-SOX17 (216-415) |
| + | + | + | GST-APC (954-1203 ) |
| - | - | + | MBP negative control |

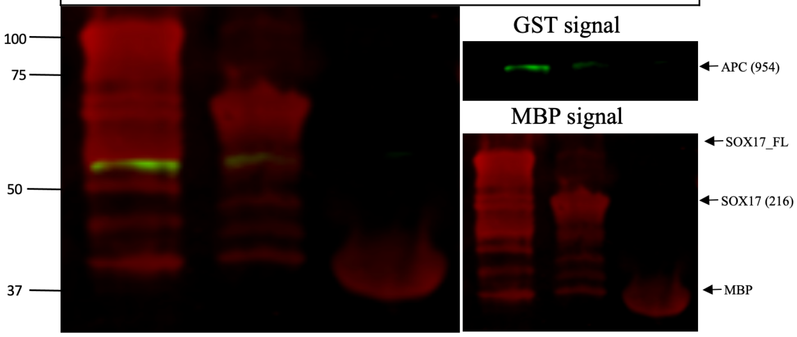


**Supplemental Figure 5: Pull-down of GST-APC$^{954-1203}$ containing the 15R region.** GST-APC$^{954-1203}$ was tested against the following MBP tagged proteins: Lane 1: SOX17$^{FL}$, Lane 2: SOX17$^{216-415}$, and Lane 3: MBP alone (negative control). Lane 1 and 2: GST-APC$^{954-1203}$ shows fragmentation represented in the gel by multiple bands below the expected target protein. Arrows point to the expected molecular weight of the target protein fragment (green signal GST; red signal MBP).

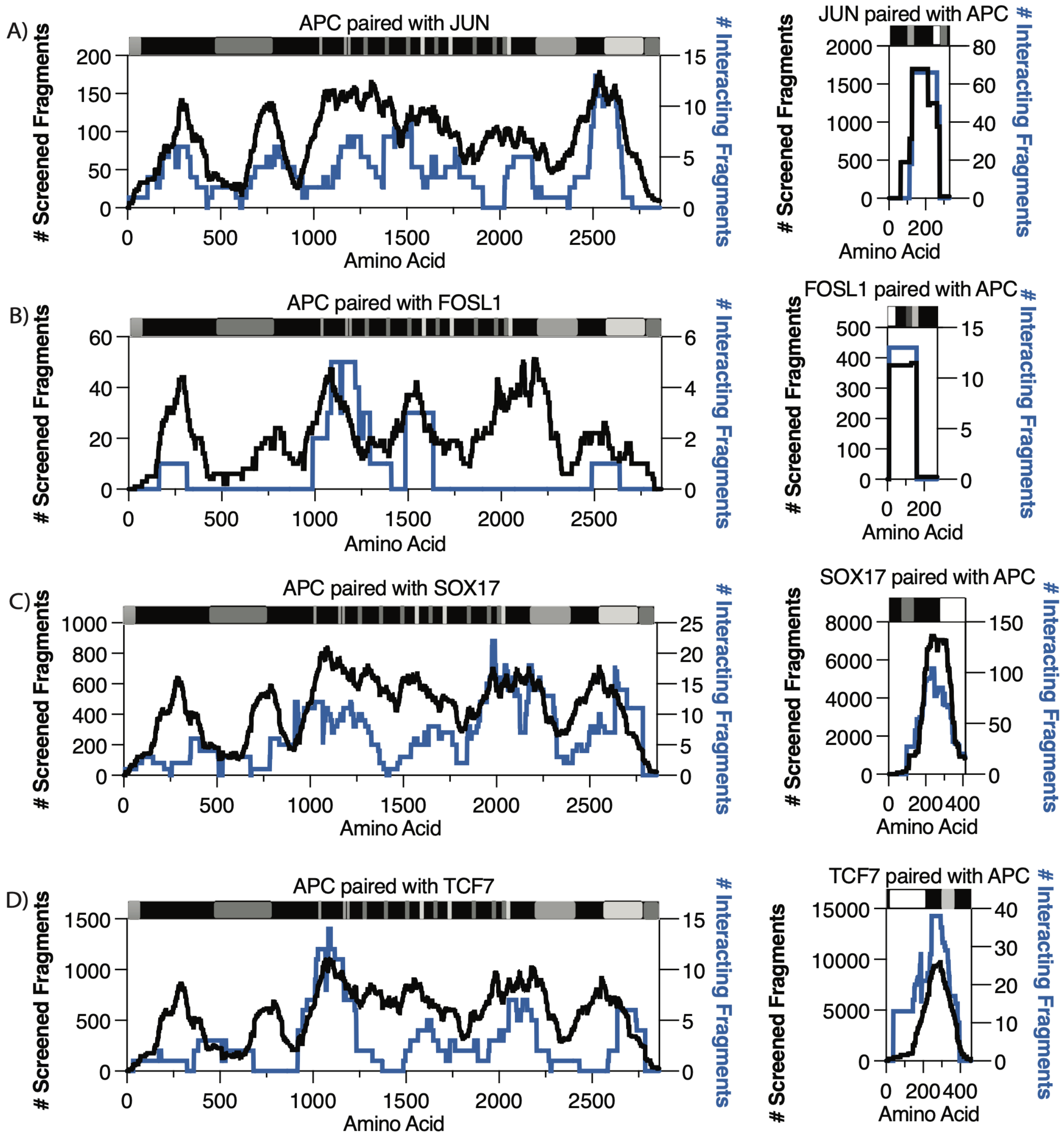


**Supplemental Figure 6: High-resolution interaction mapping of APC with Transcription factors. A, B, C, D)** APC domain are marked in order: oligomerization 6-75 aa.; armadillo 453-767 aa.; 15R repeat (amino acids 1020-1034; 11555-1169; 1172-1186); 20R repeats (amino acids 1260-1280; 1372-1393; 1486-1509; 1637-1660; 1841-1865; 1950-1972) seven domains *dark grey; SAMP 1-3 repeats (amino acids 1567-1588; 1717-1737; 2031-2051) three domains *light grey; Basic domain 2224-2575 aa.; EB1 2670-2843 aa.; DLG 2772-2843 aa. **A)** JUN domain: Transactivation domain 31-59 aa.; CTNNB1 binding region DBD 252-279 aa.; leucine zipper 280-308. **B)** FOSL1 domain: CTNNB1 binding region 1-54 aa.; DBD and Leucine zipper: 165-218 aa. **C)** SOX17 domains are marked in order: HMG box 68-136 aa.; CTNNB1 binding 280-413 aa. **D)** TCF7 domain: CTNNB1 binding region 20-212 aa.; followed by HMG box 300-370 aa. (CTNNB1 binding with transcription factors: JUN, FOSL1, TCF7, and SOX17, is marked by a white box domain). Y-axis represents the total screened fragments (left) and the number of interacting fragments (right). Black traces represent total screened fragments (coverage), while blue traces represent the number of interacting fragments. The x-axis represents protein length in amino acids.

### Supplemental Tables:

**Supplemental Table 1:** List of the 60 Wnt pathway clones purchased from Genscript.

| | |
|---|---|
| **1** | Clone ID: OHu103088D<br>ORF Clones (Accession No.): NM_001354896.1(ORF Sequence),8,583 bp ,<br>Vector: pcDNA3.1+/C-(K)-DYK |
| **2** | Clone ID: OHu61694D<br>ORF Clones (Accession No.): XM_011522682.2(ORF Sequence),2,748 bp ,<br>Vector: pcDNA3.1+/C-(K)-DYK |
| **3** | Clone ID: OHu26644D<br>ORF Clones (Accession No.): XM_017025192.1(ORF Sequence),2,529 bp ,<br>Vector: pcDNA3.1+/C-(K)-DYK |
| **4** | Clone ID: OHu29289D<br>ORF Clones (Accession No.): NM_012342.2(ORF Sequence),780 bp ,<br>Vector: pcDNA3.1+/C-(K)-DYK |
| **5** | Clone ID: OHu29786D<br>ORF Clones (Accession No.): NM_005454.2(ORF Sequence),801 bp ,<br>Vector: pcDNA3.1+/C-(K)-DYK |
| **6** | Clone ID: OHu24236D<br>ORF Clones (Accession No.): NM_004380.2(ORF Sequence),7,326 bp ,<br>Vector: pcDNA3.1+/C-(K)-DYK |
| **7** | Clone ID: OHu17197D<br>ORF Clones (Accession No.): XM_017005738.1(ORF Sequence),2,343 bp ,<br>Vector: pcDNA3.1+/C-(K)-DYK |
| **8** | Clone ID: OHu08802D<br>ORF Clones (Accession No.): NM_022802.2(ORF Sequence),2,955 bp ,<br>Vector: pcDNA3.1+/C-(K)-DYK |
| **9** | Clone ID: OHu38304D<br>ORF Clones (Accession No.): XM_005272261.3(ORF Sequence),1,323 bp ,<br>Vector: pcDNA3.1+/C-(K)-DYK |
| **10** | Clone ID: OHu27309D<br>ORF Clones (Accession No.): XM_017001855.1(ORF Sequence),243 bp ,<br>Vector: pcDNA3.1+/C-(K)-DYK |
| **11** | Clone ID: OHu26635D<br>ORF Clones (Accession No.): XM_011529176.1(ORF Sequence),765 bp ,<br>Vector: pcDNA3.1+/C-(K)-DYK |
| **12** | Clone ID: OHu17439D<br>ORF Clones (Accession No.): XM_011516632.2(ORF Sequence),2,328 bp ,<br>Vector: pcDNA3.1+/C-(K)-DYK |
| **13** | Clone ID: OHu12318D<br>ORF Clones (Accession No.): XM_017008652.1(ORF Sequence),1,101 bp ,<br>Vector: pcDNA3.1+/C-(K)-DYK |
| **14** | Clone ID: OHu10983D<br>ORF Clones (Accession No.): NM_005438.4(ORF Sequence),813 bp ,<br>Vector: pcDNA3.1+/C-(K)-DYK |

| | |
|---|---|
| **15** | Clone ID: OHu15341D<br>ORF Clones (Accession No.): NM_145866.1(ORF Sequence),1,998 bp ,<br>Vector: pcDNA3.1+/C-(K)-DYK |
| **16** | Clone ID: OHu23496D<br>ORF Clones (Accession No.): NM_003506.3(ORF Sequence),2,118 bp ,<br>Vector: pcDNA3.1+/C-(K)-DYK |
| **17** | Clone ID: OHu23207D<br>ORF Clones (Accession No.): NM_002093.3(ORF Sequence),1,299 bp ,<br>Vector: pcDNA3.1+/C-(K)-DYK |
| **18** | Clone ID: OHu23027D<br>ORF Clones (Accession No.): NM_016269.4(ORF Sequence),1,197 bp ,<br>Vector: pcDNA3.1+/C-(K)-DYK |
| **19** | Clone ID: OHu11817D<br>ORF Clones (Accession No.): NM_145203.5(ORF Sequence),1,011 bp ,<br>Vector: pcDNA3.1+/C-(K)-DYK |
| **20** | Clone ID: OHu00214D<br>ORF Clones (Accession No.): NM_001256686.1(ORF Sequence),1,173 bp ,<br>Vector: pcDNA3.1+/C-(K)-DYK |
| **21** | Clone ID: OHu10985D<br>ORF Clones (Accession No.): NM_152221.2(ORF Sequence),1,248 bp ,<br>Vector: pcDNA3.1+/C-(K)-DYK |
| **22** | Clone ID: OHu49264D<br>ORF Clones (Accession No.): XM_005255800.3(ORF Sequence),897 bp ,<br>Vector: pcDNA3.1+/C-(K)-DYK |
| **23** | Clone ID: OHu11688D<br>ORF Clones (Accession No.): NM_001320.6(ORF Sequence),645 bp ,<br>Vector: pcDNA3.1+/C-(K)-DYK |
| **24** | Clone ID: OHu23777D<br>ORF Clones (Accession No.): NM_012242.3(ORF Sequence),798 bp ,<br>Vector: pcDNA3.1+/C-(K)-DYK |
| **25** | Clone ID: OHu31152D<br>ORF Clones (Accession No.): NM_014421.2(ORF Sequence),777 bp ,<br>Vector: pcDNA3.1+/C-(K)-DYK |
| **26** | Clone ID: OHu29690D<br>ORF Clones (Accession No.): XM_017013316.1(ORF Sequence),672 bp ,<br>Vector: pcDNA3.1+/C-(K)-DYK |
| **27** | Clone ID: OHu33084D<br>ORF Clones (Accession No.): NM_001330311.1(ORF Sequence),2,085 bp ,<br>Vector: pcDNA3.1+/C-(K)-DYK |
| **28** | Clone ID: OHu18501D<br>ORF Clones (Accession No.): NM_004422.2(ORF Sequence),2,208 bp ,<br>Vector: pcDNA3.1+/C-(K)-DYK |
| **29** | Clone ID: OHu37748D<br>ORF Clones (Accession No.): XM_005247172.2(ORF Sequence),2,154 bp ,<br>Vector: pcDNA3.1+/C-(K)-DYK |

| | |
|---|---|
| **30** | Clone ID: OHu24407D<br>ORF Clones (Accession No.): NM_005479.3(ORF Sequence),837 bp ,<br>Vector: pcDNA3.1+/C-(K)-DYK |
| **31** | Clone ID: OHu01956D<br>ORF Clones (Accession No.): NM_012083.2(ORF Sequence),699 bp ,<br>Vector: pcDNA3.1+/C-(K)-DYK |
| **32** | Clone ID: OHu13313D<br>ORF Clones (Accession No.): NM_003505.1(ORF Sequence),1,941 bp ,<br>Vector: pcDNA3.1+/C-(K)-DYK |
| **33** | Clone ID: OHu25356D<br>ORF Clones (Accession No.): NM_002228.3(ORF Sequence),993 bp ,<br>Vector: pcDNA3.1+/C-(K)-DYK |
| **34** | Clone ID: OHu58778D<br>ORF Clones (Accession No.): XM_011545029.1(ORF Sequence),4,986 bp ,<br>Vector: pcDNA3.1+/C-(K)-DYK |
| **35** | Clone ID: OHu98210D<br>ORF Clones (Accession No.): XM_017013844.1(ORF Sequence),1,410 bp ,<br>Vector: pcDNA3.1+/C-(K)-DYK |
| **36** | Clone ID: OHu07491D<br>ORF Clones (Accession No.): XM_011523537.1(ORF Sequence),1,488 bp ,<br>Vector: pcDNA3.1+/C-(K)-DYK |
| **37** | Clone ID: OHu99426D<br>ORF Clones (Accession No.): XM_017029735.1(ORF Sequence),1,722 bp ,<br>Vector: pcDNA3.1+/C-(K)-DYK |
| **38** | Clone ID: OHu27910D<br>ORF Clones (Accession No.): XM_011536974.1(ORF Sequence),1,389 bp ,<br>Vector: pcDNA3.1+/C-(K)-DYK |
| **39** | Clone ID: OHu27909D<br>ORF Clones (Accession No.): XM_011536972.2(ORF Sequence),1,401 bp ,<br>Vector: pcDNA3.1+/C-(K)-DYK |
| **40** | Clone ID: OHu25308D<br>ORF Clones (Accession No.): NM_021627.2(ORF Sequence),1,767 bp ,<br>Vector: pcDNA3.1+/C-(K)-DYK |
| **41** | Clone ID: OHu18617D<br>ORF Clones (Accession No.): NM_002615.5(ORF Sequence),1,254 bp ,<br>Vector: pcDNA3.1+/C-(K)-DYK |
| **42** | Clone ID: OHu27073D<br>ORF Clones (Accession No.): NM_003012.4(ORF Sequence),942 bp ,<br>Vector: pcDNA3.1+/C-(K)-DYK |
| **43** | Clone ID: OHu17742D<br>ORF Clones (Accession No.): NM_025237.2(ORF Sequence),639 bp ,<br>Vector: pcDNA3.1+/C-(K)-DYK |
| **44** | Clone ID: OHu24692D<br>ORF Clones (Accession No.): NM_022454.3(ORF Sequence),1,242 bp ,<br>Vector: pcDNA3.1+/C-(K)-DYK |

| | |
|---|---|
| **45** | Clone ID: OHu09015D<br>ORF Clones (Accession No.): XM_017007185.1(ORF Sequence),1,542 bp ,<br>Vector: pcDNA3.1+/C-(K)-DYK |
| **46** | Clone ID: OHu39688D<br>ORF Clones (Accession No.): XM_006714678.3(ORF Sequence),1,380 bp ,<br>Vector: pcDNA3.1+/C-(K)-DYK |
| **47** | Clone ID: OHu10988D<br>ORF Clones (Accession No.): NM_138959.2(ORF Sequence),1,572 bp ,<br>Vector: pcDNA3.1+/C-(K)-DYK |
| **48** | Clone ID: OHu23083D<br>ORF Clones (Accession No.): XM_011509804.1(ORF Sequence),1,563 bp ,<br>Vector: pcDNA3.1+/C-(K)-DYK |
| **49** | Clone ID: OHu14993D<br>ORF Clones (Accession No.): NM_007191.4(ORF Sequence),1,137 bp ,<br>Vector: pcDNA3.1+/C-(K)-DYK |
| **50** | Clone ID: OHu22106D<br>ORF Clones (Accession No.): NM_005430.3(ORF Sequence),1,110 bp ,<br>Vector: pcDNA3.1+/C-(K)-DYK |
| **51** | Clone ID: OHu18435D<br>ORF Clones (Accession No.): NM_003394.3(ORF Sequence),1,167 bp ,<br>Vector: pcDNA3.1+/C-(K)-DYK |
| **52** | Clone ID: OHu11384D<br>ORF Clones (Accession No.): NM_003391.2(ORF Sequence),1,080 bp ,<br>Vector: pcDNA3.1+/C-(K)-DYK |
| **53** | Clone ID: OHu23758D<br>ORF Clones (Accession No.): NM_030753.4(ORF Sequence),1,065 bp ,<br>Vector: pcDNA3.1+/C-(K)-DYK |
| **54** | Clone ID: OHu74051D<br>ORF Clones (Accession No.): XM_011544319.2(ORF Sequence),1,338 bp ,<br>Vector: pcDNA3.1+/C-(K)-DYK |
| **55** | Clone ID: OHu68294D<br>ORF Clones (Accession No.): XM_011541597.2(ORF Sequence),1,119 bp ,<br>Vector: pcDNA3.1+/C-(K)-DYK |
| **56** | Clone ID: OHu94538D<br>ORF Clones (Accession No.): XM_017007127.1(ORF Sequence),1,182 bp ,<br>Vector: pcDNA3.1+/C-(K)-DYK |
| **57** | Clone ID: OHu05697D<br>ORF Clones (Accession No.): NM_006522.3(ORF Sequence),1,095 bp ,<br>Vector: pcDNA3.1+/C-(K)-DYK |
| **58** | Clone ID: OHu18078D<br>ORF Clones (Accession No.): NM_004625.3(ORF Sequence),1,047 bp ,<br>Vector: pcDNA3.1+/C-(K)-DYK |
| **59** | Clone ID: OHu96059D<br>ORF Clones (Accession No.): XM_011543625.2(ORF Sequence),1,011 bp ,<br>Vector: pcDNA3.1+/C-(K)-DYK |

| | |
|---|---|
| **60** | Clone ID: OHu17830D<br>ORF Clones (Accession No.): NM_003396.2(ORF Sequence),1,071 bp ,<br>Vector: pcDNA3.1+/C-(K)-DYK |

**Supplemental Table 2: A list of stringent known protein-protein interactions recovered.** Around 50% of the known interactions (37 out of 74) were detected in both orientations (protein fragments being associated with AD "Orient 1" and associated with DBD "Orient 2"). Columns 1 and 2 list Protein 1 and Protein 2, which are the tested protein pair. The following two columns, Orient 1 and Orient 2 show how many times the protein pair is detected in each orientation (Orient 1 is AD-associated; Orient 2 is DBD-associated) and then followed by a 3-AT competitive inhibitor condition to determine the number of pairs detected in 2 mM vs. 5 mM. Significant interactions filtered by $Log_2FCmax$ and FDRmin values. The number of libraries shows if the pairs are captured in a single library or both (with 2 being the maximum). The unique fragment pairs represent the number of unique fragments captured for each protein pair. The APID concludes its known interaction = 1.

| Protein1 | Protein2 | Orient1 | Orient2 | 2mM | 5mM | $log_2FCmax$ | FDRmin | Library | Unique Pairs | APID |
|---|---|---|---|---|---|---|---|---|---|---|
| AXIN1 | APC | 227 | 64 | 159 | 132 | 8.82043272 | 0 | 2 | 201 | 1 |
| DVL3 | DVL1 | 18 | 8 | 14 | 12 | 7.27107565 | 2.56E-231 | 2 | 17 | 1 |
| CTBP1 | CTBP2 | 3 | 8 | 8 | 3 | 8.94301438 | 5.75E-169 | 2 | 9 | 1 |
| WIF1 | WNT7A | 3 | 7 | 7 | 3 | 7.98946273 | 7.72E-121 | 2 | 6 | 1 |
| APC | AXIN2 | 30 | 122 | 77 | 75 | 7.52024269 | 2.10E-97 | 2 | 123 | 1 |
| AXIN2 | AXIN1 | 49 | 13 | 34 | 28 | 5.50507240 | 3.16E-92 | 2 | 52 | 1 |
| CTBP1 | APC | 73 | 5 | 46 | 32 | 6.19893196 | 1.52E-21 | 2 | 67 | 1 |
| AXIN2 | CTNNB1 | 9 | 2 | 9 | 2 | 6.05062835 | 6.77E-20 | 2 | 9 | 1 |
| CREBBP | APC | 280 | 79 | 147 | 212 | 5.57788281 | 3.70E-17 | 2 | 307 | 1 |
| CTNNB1 | CTBP2 | 5 | 14 | 8 | 11 | 3.25745651 | 1.33E-10 | 2 | 19 | 1 |
| DVL3 | CSNK2A1 | 1 | 2 | 2 | 1 | 2.95446117 | 2.11E-10 | 2 | 3 | 1 |
| AXIN1 | LRP5 | 11 | 38 | 27 | 22 | 5.18518182 | 2.92E-10 | 2 | 40 | 1 |
| DVL1 | APC | 50 | 10 | 31 | 29 | 4.15410934 | 1.12E-09 | 2 | 55 | 1 |
| DVL3 | AXIN1 | 30 | 21 | 19 | 32 | 5.55985323 | 2.94E-09 | 2 | 42 | 1 |
| CTNNB1 | AXIN1 | 7 | 6 | 8 | 5 | 5.02547308 | 3.20E-09 | 2 | 12 | 1 |
| AXIN1 | DVL1 | 10 | 11 | 11 | 10 | 5.63935026 | 5.39E-07 | 2 | 17 | 1 |
| WNT3A | WNT1 | 3 | 3 | 2 | 4 | 3.15960247 | 5.42E-07 | 1 | 5 | 1 |
| SOST | LRP5 | 2 | 5 | 3 | 4 | 2.61087125 | 5.45E-07 | 2 | 7 | 1 |
| CTNNB1 | CREBBP | 12 | 2 | 8 | 6 | 3.03869564 | 6.17E-07 | 2 | 14 | 1 |
| FZD1 | WNT3A | 4 | 4 | 3 | 5 | 6.09936507 | 9.55E-07 | 2 | 7 | 1 |
| DVL2 | DVL3 | 3 | 2 | 2 | 3 | 2.44493789 | 4.54E-06 | 2 | 5 | 1 |
| CTNNB1 | SOX17 | 7 | 8 | 5 | 10 | 5.18400417 | 1.26E-05 | 2 | 12 | 1 |
| DVL2 | AXIN1 | 6 | 1 | 4 | 3 | 2.61599891 | 1.49E-05 | 2 | 7 | 1 |
| WNT1 | LRP5 | 14 | 7 | 10 | 11 | 4.97492927 | 4.38E-05 | 2 | 16 | 1 |
| CSNK2A | CREBBP | 1 | 1 | 1 | 1 | 3.45739677 | 6.89E-05 | 2 | 2 | 1 |
| CTBP2 | DVL2 | 4 | 1 | 2 | 3 | 2.99606395 | 7.51E-05 | 2 | 4 | 1 |
| SENP2 | AXIN1 | 4 | 13 | 10 | 7 | 5.56907543 | 0.00010856 | 2 | 13 | 1 |

| | | | | | | | | | | |
|---|---|---|---|---|---|---|---|---|---|---|
| WNT1 | WIF1 | 9 | 1 | 3 | 7 | 3.14992036 | 0.0001673 | 2 | 9 | 1 |
| VANGL1 | DVL3 | 3 | 1 | 2 | 2 | 3.09639402 | 0.0003971 | 2 | 4 | 1 |
| DVL1 | CTNNB1 | 3 | 3 | 5 | 1 | 3.15561286 | 0.0004161 | 2 | 5 | 1 |
| WNT3A | WNT3 | 4 | 3 | 2 | 5 | 2.79734786 | 0.0004748 | 2 | 6 | 1 |
| WNT7A | WNT5A | 2 | 2 | 3 | 1 | 3.02808694 | 0.0006909 | 1 | 3 | 1 |
| DVL3 | CTNNB1 | 4 | 4 | 5 | 3 | 3.96613970 | 0.0010586 | 2 | 8 | 1 |
| WNT3 | FZD1 | 4 | 1 | 2 | 3 | 2.81715288 | 0.0018885 | 2 | 4 | 1 |
| VANGL1 | DVL1 | 2 | 3 | 3 | 2 | 2.83187419 | 0.0019781 | 2 | 2 | 1 |
| TCF7 | CTNNB1 | 1 | 2 | 0 | 3 | 2.06717158 | 0.0072836 | 1 | 3 | 1 |

**Supplemental Table 3: A list of tested fragments used for pull-down.** TWIST fragments: APC, FOSL1, JUN, and TCF7. While SOX17 fragments were PCR amplified using primers compatible with Electra cloning.

| Protein | Fragments bp location | Amino acid location |
|---|---|---|
| APC | 2815-3714 | 938-1239 |
| | 2860-3609 | 954-1203 |
| | 4534-5283 | 1512-1761 |
| | 5869-6696 | 1956-2232 |
| | 7615-8316 | 2539-2772 |
| FOSL1 | 1-846 | 1-282 |
| | 115-813 | 40-282 |
| JUN | 1-996 | 1-331 |
| | 298-996 | 100-331 |
| TCF7 | 1-1383 | 1-461 |
| | 454-1077 | 152-359 |
| SOX17 | 1-1245 | 1-415 |
| | 261-862 | 88-287 |
| | 646-1245 | 216-415 |
